# A Fibroblast State Choreographs an Epithelial YAP-dependent Regenerative Program Essential to (Pre)malignancy via ECM-mediated Mechanotransduction

**DOI:** 10.1101/2025.07.11.661192

**Authors:** Deng Pan, Philippe Gascard, Joseph A. Caruso, Chira Chen-Tanyolac, Veena Sangwan, Nicholas Bertos, Sophie Camilleri-Broet, Julie Berube, Spyridon Oikonomopoulos, Michael K. Strasser, David L. Gibbs, Joanna Bons, Jordan B. Burton, Jacob P. Rose, Samah Shah, Rosemary Bai, Stuart Lee, Daffolyn Rachael Fels-Elliott, Shoval Miyara, Uri Alon, Anatoly Urisman, Ioannis Ragoussis, Sui Huang, Birgit Schilling, Lorenzo E. Ferri, Thea D. Tlsty

## Abstract

Chronic lung injury generates metaplasia which occasionally, but ominously, progresses to squamous dysplasia and squamous lung cancer. To identify mechanisms through which disrupted tissue homeostasis contributes to malignant initiation and progression, we used *in vivo* and *in vitro* heterotypic recombinant models of human <u>b</u>ronchial <u>e</u>pithelial <u>c</u>ells (hBECs) and fibroblasts. We demonstrate that injury-associated TGF-β signaling creates a fibroblast state dependent upon HSP47 upregulation. These fibroblasts accumulated collagen, thus elevating tissue stiffness and activating mechanosignaling that sustained YAP-dependent embryonic-like, pro-malignant activities in adjacent hBECs. This <u>S</u>tress/<u>T</u>ension-Instructive <u>F</u>ibroblast (STIF) state, exhibited by stressed fibroblasts in premalignant and malignant lesions across multiple cancer types, was sufficient to reprogram disease-free hBECs to metaplasia and to drive hBECs with compromised tumor suppressor function to dysplasia, yet could be inhibited and reversed. STIFs suffice to activate epithelial phenotypes reminiscent of oncogene-mediated cell transformation and induce (pre)malignancy via increased force transmission, providing novel targets for prevention.

**Statement of significance:** Tissue injury creates a regenerative pro-tumorigenic <u>S</u>tress/<u>T</u>ension-Instructive <u>F</u>ibroblast (STIF) state which is sufficient to activate a YAP-dependent, pre-malignant program to induce or unmask pre-cancerous phenotypes in epithelial cells through mechanotransduction. Inhibition of STIF activity or mechanosignaling prevents metaplasia and progression to dysplasia.

**Highlights:**

1. Tissue injury creates a pro-tumorigenic Stress/Tension-Instructive Fibroblast (STIF) state in multiple organs that precedes and persists through cancer
2. STIF signaling alone, working through fibroblasts and not epithelial cells, is sufficient to activate embryonic-like plasticity and induce epithelial pre-cancerous metaplastic lesions
3. STIFs program (pre)malignant phenotypes in adjacent epithelial cells through mechanosignaling by activating YAP prior to tumor formation
4. Inhibiting STIFs or mechanosignaling prevents/reverts metaplasia and prevents progression to dysplasia

## Introduction

Morphologically normal epithelial cells in humans and mice can harbor mutations and copy number alterations in genes commonly associated with malignancy (TP53, PI3KCA, KRAS, and EGFR) but, paradoxically, do not commonly progress to cancer ^1–6^ Similarly, individuals can carry pre-cancerous lesions with oncogenic mutation(s) in multiple adult tissues that don’t progress to overt cancer during their lifetime^7,8^. These observations, summarized in a recent insightful review ^9^, suggest that factors beyond epithelial mutations and oncogene activation, such as stromal contributions, are needed for disease initiation and progression.

In a related paradox, many carcinogens drive the initiation and progression of tumors, but are not mutagens (i.e., chronic inflammation, obesity, or aging)^10–13^. The mechanisms underlying this association have not been clearly established but studies suggest they are non-mutational. In cancers driven by chronic inflammation, such as lung squamous cell carcinomas (LSCCs), alterations in TP53 and CDKN2A are common; however, unlike lung adenocarcinoma, LSCCs rarely exhibit typical oncogenic drivers, such as EGFR, KRAS, or ALK fusions^14,15^ demonstrating that cancer can develop in the absence of these well-studied drivers, with chronic inflammation potentially serving as an effective substitute.

These paradoxes can be reconciled by considering the key role of the microenvironment in carcinogenesis. Previous and current studies have challenged the traditional epithelial/mutational-centric view of the genesis of cancer and demonstrated that stromal signaling is a critical aspect of epithelial cell behavior in development and disease ^15–17^. Stromal signals can induce normal cells to become cancer precursors and non-tumorigenic immortalized cells to become malignant ^16–19^. Conversely, stromal signals can also repress malignant phenotypes when tumor cells are placed in a “normalizing” microenvironment both *in vitro* and *in vivo* ^20–25^.

Here, we address the role of stromal components, specifically fibroblasts, in inducing and driving (pre)malignant phenotypes. Historically, it has been difficult to study cancer initiation since it is often sporadic in location and time. Instructively, areas of chronic injury exhibit a higher probability of generating malignancies ^13,26,27^. One example is LSCC, a subtype of non-small-cell lung cancer, accounting for 30-35% of lung cancer cases ^28,29^. Chronic tissue damage caused by smoking and/or air pollution triggers subsequent chronic inflammation and injury, which often leads to an adaptive and wound-healing, regenerative tissue response known as metaplasia, where the original differentiated cell type is replaced by another mature, differentiated cell type that is not originally present ^26,27^. In the bronchial airway, pseudostratified ciliated and secretory bronchial epithelial cells are replaced by squamous epithelial cells, which increase tissue adaptiveness under injurious conditions. Although protective, metaplasia is associated with an increased risk of progressing to dysplasia and cancer with the underlying mechanisms still not well understood ^28,30,31^. Hence, chronic inflammation activates a change in cell identity and generates a metaplastic lesion (squamous metaplasia) that is directly linked to an increased incidence of a specific cancer type (squamous lung cancer), spotlighting the temporal and spatial origins of malignant potential and facilitating the study of early malignant progression. Leveraging this insight, we focused our study on the earliest steps in the generation of LSCC. We demonstrate that increased collagen deposition by fibroblasts exposed to stress signals, driven by upregulation of HSP47, resulted in increased stromal stiffness and activation of mechanosignaling that induces sustained YAP signaling in epithelial cells. The ensuing sustained activation of embryonal-like plasticity during attempted regeneration of the chronically injured tissue, prior to tumor formation, results in expression of metaplastic and dysplastic phenotypes in overlying epithelial cells. Notably, inhibition of HSP47 activity or mechanosignaling prevents and reverts formation of metaplasia and prevents dysplasia, demonstrating that activation of pro-malignant fibroblast signaling in chronically injured tissues and premalignant lesions (developing prior to tumor formation), can be modulated for clinical prevention of disease.

## Results

### Fibroblasts Experiencing Oxidative Stress are Sufficient to Induce Squamous Metaplasia when Co-cultured with Primary hBECs

To test the role of stromal signaling, specifically fibroblasts, in LSCC initiation and progression, we used human tissue recombinants in optimized *in vitro* co-culture and murine *in vivo* models (Figure 1A). We assessed epithelial cell characteristics including differentiation, proliferation, cell type composition, activation of embryonal-like plasticity, genomic stability and pathological responses (Figures 1 and S1 – S3). We focused on fibroblasts because our previous studies identifying and characterizing Carcinoma-Associated Fibroblasts (CAFs)^18^ indicated that fibroblasts play a prominent role in orchestrating epithelial responses in malignancy. Since LSCC has been reported to arise from bronchial (small airway) epithelial cells (hBECs), we used patient-derived human bronchial organoid cultures enriched in airway stem cells, as previously described ^32^. Our optimized Air-Liquid Interface (ALI) culture model ^33^ mimicked the physiological differentiation of airway epithelium more faithfully than an organoid model (Figure S1B,D,E). In-depth characterization demonstrated that hBECs gave rise to ALI cultures with functional, well-differentiated and polarized bronchial epithelial structures resembling the human airway epithelium *in vivo* (Figure S1A-G). The observations obtained with the ALI cultures were recapitulated in an *in vivo* setting, where we optimized the growth of primary hBECs in murine hosts (Figure S1H) within a complex *in vivo* environment containing multiple cell types and ECM components.

**Figure 1:**
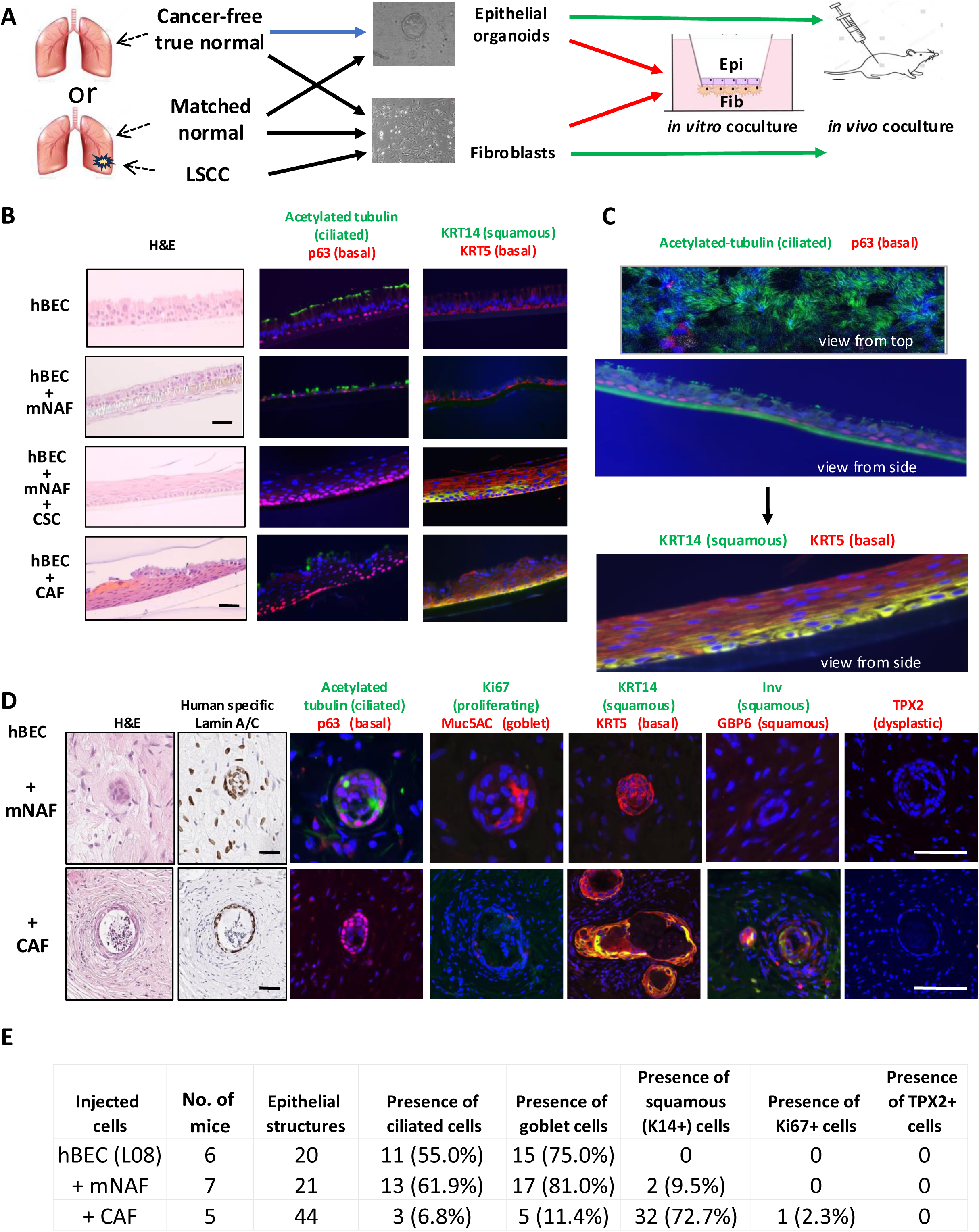
Fibroblasts Experiencing Oxidative Stress are Sufficient to Induce Squamous Metaplasia when Co-cultured with Primary hBECs. (**A**) Bronchial epithelial and stromal cell co-culture flowchart *in vitro* and *in vivo*. (B) hBECs cocultured with mNAFs were treated with vehicle control or 50ng/ml of CSC for 21 days in ALI coculture. hBECs alone in ALI were used as a control. Epithelium was assessed for morphology by H&E staining and cell differentiation by immunostaining for bronchial epithelial and squamous metaplastic markers. (C) Representative IHC of hBECs co-cultured with mNAF (top) or CAF (bottom) showing epithelial structures expressing bronchial epithelial markers (top) or squamous markers (bottom). (D) Bronchial epithelial cells were co-injected subcutaneously with mNAFs or CAFs into immunocompromised NSG mice. After four weeks, tissue structures were assessed for morphology by H&E staining and stained for human-specific Lamin A/C to localize the formed human epithelial structures. Bronchial epithelial cell differentiation was assessed by immunostaining for bronchial epithelial and squamous metaplasia markers. (E) Human epithelial structures formed in mice were quantified according to their differentiation status based on marker expression profile.

To focus on the mechanistic insights derived from our previous CAF studies, we co-cultured hBECs with fibroblasts isolated from either healthy or malignant human lung tissue from multiple, treatment-naïve patients diagnosed with LSCC (Materials and Methods; Figure 1A and Table S1). Fibroblasts from airway distal to the cancer (matched Normal-Associated Fibroblasts; mNAFs) were ultimately compared to fibroblasts obtained from within the tumor tissue, CAFs, both identities confirmed by immunofluorescent staining (Figure S2B). After 21 or 42 days of co-culture, mNAFs maintained an organized pseudostratified epithelial architecture with minimal proliferation and proper expression of apical bronchial and basal cell differentiation markers (Figures 1B-C and S2A). As expected, there was no obvious expression of cytokeratin 14 (KRT14), a marker of squamous metaplasia (Figure 1B). Fibroblasts generated a thin layer of basement membrane and a thick layer of collagens beneath hBECs in the co-cultured structure (Figure S2D) which enabled cell-cell interactions through both soluble factors and ECM deposition due to the close contact between the two cell types. Indeed, we documented that ECM proteins, such as collagens, extended from the lower fibroblast layer to the top epithelial layer through the pores of the transwell membrane and that the quality of collagen deposition differed between mNAFs and CAFs (Figure S2D and F).

Cigarette smoke and cigarette smoke condensate (CSC) cause oxidative stress (increased production of ROS) and are known risk factors for lung cancer^34–36^. Exposure of the ALI hBEC + mNAF recombinant co-cultures to CSC resulted in a dramatic appearance of metaplastic squamous epithelial cells replacing the ciliated-secretory epithelial cells described previously. The loss of ciliated and secretory cells was accompanied by basal cell hyperplasia and robust expression of squamous markers, cytokeratin 14 (KRT14) and the basal marker cytokeratin 5 (KRT5) (Figure 1B), identical to that reported *in vivo* in injured human lungs. Importantly, we established that this squamous identity exhibited by recombinant ALI metaplastic structures recapitulated an esophageal squamous cell identity, as observed *in vivo* during LSCC progression (Figure S3). Esophageal and lung tissue share a common embryonic progenitor ^37,38^.

CAFs also exist in a state of oxidative stress^39,40^. Additionally, previous studies with human cell recombinants had demonstrated that co-culture of CAFs or CAF-like fibroblasts with disease-free prostatic^18^ or breast epithelial cells^41^ generated precancerous lesions. When hBECs were co-cultured with CAFs focal areas of epithelium adopted a metaplastic squamous (precancerous) morphology with expression of KRT14 and involucrin (INV). (Figures 1B-C – lower panels, S2A and S2E). Entire transwell membranes containing the epithelium co-cultured with either mNAFs or CAFs were sectioned to assess the presence and extent of squamous epithelial metaplasia (Figure S2G). Whereas there was no evidence of squamous epithelium in ALI co-cultures with mNAFs (Figure S2G), in contrast, over 20% of the bronchial epithelial surface area was replaced by squamous epithelium (metaplasia) after 21 days in co-culture with CAFs (Figure 1B-C, S2A and S2G) and 100% after 42 days (Figure 1C, bottom panel and S2G).

The hBECs above were derived from matched histologically normal tissues distal to the LSCC growth but in the same patient. Studies reported that histologically normal tissues adjacent to tumors may be exposed to stress signals released from the tumor and may differ from cells isolated from healthy tissues ^42^. To rule out the possibility that the utilized hBECs might have been “primed” by a “stressed” tumor microenvironment (even from a distance), we investigated the response of “true” normal bronchial organoids (true-hBECs) and trueNFs obtained from disease-free autopsy lung samples. As expected, true-hBECs recapitulated results obtained with hBECs used above (Figure S4A). Additionally, trueNFs also recapitulated the above described mNAF results supporting homeostatic differentiation of true-hBECs without induction of any metaplastic (squamous) response (Figure S4A).

Finally, to extend the ALI culture-based observations of CAFs’ ability to induce squamous metaplasia to an *in vivo* context, hBECs were co-injected subcutaneously with different types of fibroblasts into immunocompromised mice. After four weeks, we confirmed outgrowths of human-specific epithelial structures. In accordance with our previous *in vitro* ALI observations, hBECs co-injected with mNAFs gave rise to human lung bronchial epithelium and those co-injected with CAFs exhibited squamous metaplastic phenotypes (over 70% of the structures) with no notable increase in the population of Ki67-positive proliferative cells across all the conditions (Figure 1D-E). These *in vivo* findings were consistent with our *in vitro* ALI observations and the histological morphology of patient samples *in vivo* as they progress from healthy tissue to metaplasia, dysplasia and tumor (Figure S1A), further supporting the notion that fibroblasts experiencing oxidative stress were sufficient to induce squamous metaplasia.

### ECM Remodeling is Necessary and Sufficient to Induce Squamous Metaplasia in hBECs

As fibroblasts exert many of their effects through secretion of soluble factors and/or remodeling of the ECM, we investigated whether soluble factors released by fibroblasts could induce squamous metaplasia. We employed a single-sided, indirect co-culture where hBECs and fibroblasts were cultured in physically separated compartments (Figure 2A, left) and communicated solely through soluble factors present in medium. Intriguingly, under these conditions, CAFs were unable to induce squamous metaplasia. Additionally, no discernible difference was observed in bronchial epithelial cell differentiation when hBECs were cultured with mNAFs or CAFs; ciliated and secretory cells were formed in both co-cultures (Figure 2A, left panel and histogram).

**Figure 2.**
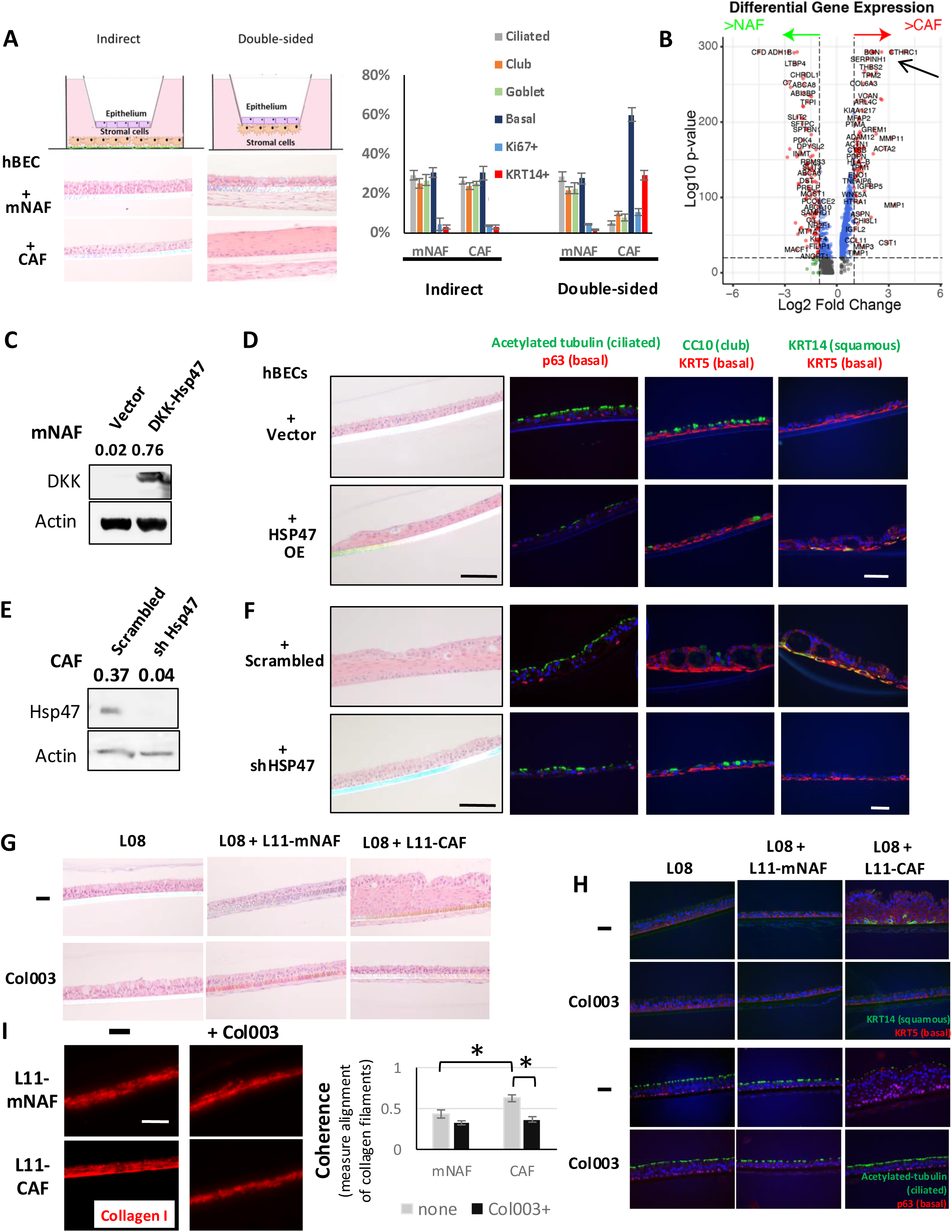
CAF-derived ECM can Induce Squamous Metaplasia in hBECs through an Hsp47-dependent Mechanism. **(A)** Schematic representation (left top) for hBECs co-cultured with mNAFs or CAFs under single-sided or double-sided conditions for 21 days. Histological assessment of co-cultures used H&E staining (left bottom). Percentages for each cell lineage generated from single-sided or double-sided cocultures quantified by immunostaining (right). **(B)** Differentially expressed genes in CAFs and mNAFs identified from scRNA-seq dataset. SERPINH1/HSP47 (arrow) was a top upregulated candidate in CAFs vs mNAFs. **(C)** A lentiviral construct encoding DKK-tagged HSP47 was introduced into mNAFs and expression assessed by western blotting using a DKK antibody. **(D)** H&E images showing bronchial epithelial cell structures co-cultured with transduced mNAFs or their controls. Immunofluorescence images showing expression of squamous and bronchial epithelial markers in hBECs. **(E)** A lentiviral construct encoding a silencing shRNA against HSP47 or scrambled shRNA was introduced into CAFs and expression was assessed by Western blotting. **(F)** Bronchial epithelial cell structures co-cultured with transduced CAFs or their controls were analyzed as in (**D**). **(G-H)** hBECs were cultured in ALI or co-cultured with mNAFs or CAFs in presence of the HSP47 inhibitor Col003 (10 μM) or vehicle control for 21 days (**G**). Resulting structures were processed as in **(F)**. **(I)** mNAFs or CAFs co-cultured in presence or absence of Col003 (10 μM) were stained for Collagen I (left). Coherence of collagen filament alignment quantified using OrientationJ in ImageJ (right).

We therefore hypothesized that the induction of metaplasia may involve ECM deposition/remodeling. Indeed, culture of hBECs on decellularized ECM obtained from CAFs induced squamous marker expression while culture on decellularized ECM from mNAFs did not (Figure S5A). To identify potential candidate proteins, we analyzed scRNA-seq data from a LSCC cohort of 21 treatment-naive patients, focusing on differentially expressed genes in CAFs compared to mNAFs. These included many ECM proteins and collagen regulators indicating that CAFs played a major role in ECM remodeling (Figures 2B and S5B-D). The identified genes were then contrasted at the proteomic level with matrisomal proteins differentially expressed in LSCC compared to matched adjacent normal lung tissues in the same cohort (Figure S5E). This analysis extended our recently published comparative findings, which identified SERPINH1, also known as Heat Shock Protein 47 (HSP47), as a significantly increased matrisomal top candidate protein in LSCC ^43^.

Several lines of evidence suggest that HSP47 activity in CAFs is an excellent candidate for the induction of metaplasia. Firstly, HSP47 is a chaperone essential for collagen synthesis, deposition, and proper assembly, ensuring the formation of a collagen network in the extracellular space. Through its involvement in collagen accumulation, this protein contributes to tissue stiffness and repair during wound healing ^44,45^. Secondly, we documented high HSP47 expression in stromal cells adjacent to LSCC, whereas its expression in matched normal tissues was barely detectable (Figure S5B)^43^. The stromal cells expressing HSP47 displayed a fibroblast-like morphology and expressed the fibroblast marker PDGFRα, confirming that fibroblasts were a major source of HSP47 in tumors (Figures S2B-C). Thirdly, we confirmed a high level of HSP47 expression in the primary LSCC samples as well as many of the cultured CAFs derived from those same samples, while HSP47 expression was low in mNAFs generated from histologically normal tissues adjacent to LSCC (Figure S5F-G). Notably, the low expression of HSP47 in mNAFs was accompanied by negligible induction of metaplasia. Finally, we observed dramatic differences in the extent of HSP47 expression in CAFs from different patients and noted that HSP47 expression in CAFs correlated with their potential to induce squamous metaplasia in hBECs (Figure S5H and Table S1). Thus, CAFs from patient L11 (expressing high levels of HSP47 *in vivo* and *in vitro*) were more potent inducers of squamous metaplasia than those from patients L09 or L10 (expressing lower levels of HSP47 *in vivo* and *in vitro*) (Figure S5E-H). These differences between CAFs may be related to patient-to-patient variability or sample collection in the vicinity of a lesion. This variability was unlikely to result from a selection in cell culture since differences in the amount of HSP47 expression in cultured cells *in vitro* were also observed *in vivo* in the corresponding tumors from which the cells were isolated (compare Figure S5E-G for lung tumors L09-L11). We also confirmed an increase in the expression of HSP47 *in situ* in CAFs neighboring areas of the transwell membrane, with a robust squamous metaplastic response in hBECs (Figures S2C and S5C). Additionally, the collagen filaments produced by CAFs were more aligned than those produced by mNAFs (Figure S2F and Figure 2I). These observations are consistent with the role of HSP47 in organizing collagen networks and with well-documented differences in collagen quality between healthy and malignant tissues ^46,47^. Taken together, these observations brought credence to our hypothesis that HSP47 might participate in CAF-induced squamous metaplasia through ECM deposition and remodeling.

The inter-patient variability provided us with a unique opportunity to identify the underlying mechanisms of CAF-induced squamous metaplasia. To determine whether HSP47 played a mechanistic role in CAF-induced squamous metaplasia, we genetically modulated HSP47 expression (Figures 2C-F and S5I). We overexpressed HSP47 (HSP47 OE) by introducing lentivirus cDNA encoding DKK-tagged HSP47 into mNAFs that originally expressed low levels of HSP47 and were not previously able to induce squamous metaplasia (Figure 2C). Notably, these mNAFs, now overexpressing HSP47, induced squamous metaplasia when co-cultured with hBECs (Figure 2D). Conversely, introducing a shRNA lentivirus (shHSP47) to silence HSP47 in CAFs, which originally expressed high levels of HSP47 and potently induced squamous metaplasia when co-cultured with hBECs, demonstrated the opposite effect (Figure 2E-F). CAFs with silenced HSP47 (Figures 2E and S5I) failed to induce squamous metaplasia, whereas apical bronchial markers were robustly expressed (Figure 2F). Finally, pharmacological abrogation of HSP47 activity in CAF via exposure to a small molecule inhibitor, Col003 ^48^, prevented squamous metaplasia (Figures 2G-H and S5J). Both collagen quantity and coherence were diminished by inhibition by Col003 (Figure 2I). Together, these results demonstrate that the increased expression of HSP47 observed in CAFs was sufficient and necessary to trigger squamous metaplasia when co-cultured with hBECs and that inhibition of HSP47 activity prevented metaplastic induction.

### Metaplasia is Induced by CAFs from Multiple Cancer Types and Fibroblasts from Pre-cancerous Lesions

Examination of the TCGA dataset revealed that high expression of HSP47 protein in tumor stroma is observed in most tumors (Figure 3A). We verified this observation via immunohistochemistry (Figure 3B), proteomic analyses (Figure S6A-B) and mRNA expression (Figure 3C) in multiple tumor types and determined that CAF obtained from these tumors could also induce squamous metaplasia in recombinant bronchial ALI co-cultures (Figures 3D-G, S6C-D). Not only did CAFs from gastric adenocarcinoma, esophageal adenocarcinoma and breast cancer exhibit high expression of HSP47 accompanied by induced squamous metaplasia, but inhibition of HSP47 activity using Col003 ablated this induction (Figures 3D-F and S6C-D). In breast cancer subtypes, HSP47 expression correlated with tumor aggression and inversely with tumor differentiation (Figure S6B). Notably, high expression of HSP47 predicts poor prognosis in multiple cancer types (Table S2). These data demonstrated that CAFs from multiple tumors share the activation of a fundamental HSP47-dependent program that can induce tumor progression as measured by induction of metaplasia. An increase in HSP47 mRNA levels was also observed in fibroblasts obtained from pre-malignant lesions of esophageal and gastric adenocarcinomas (Figures 3C) and DCIS lesions in human breast tissue (data not shown). Each exhibited the ability to induce squamous metaplasia in our recombinant co-culture ALI model (Figure 3D-G). Finally, immunohistochemistry of lung tissues at higher risk for developing lung cancer mimics the colocalization of markers seen in the ALI co-cultures (compare Figure S6F with S6G and S6H). Notably, HSP47 expression is barely detected in healthy lung bronchial tissues (Figure S6I).

**Figure 3:**
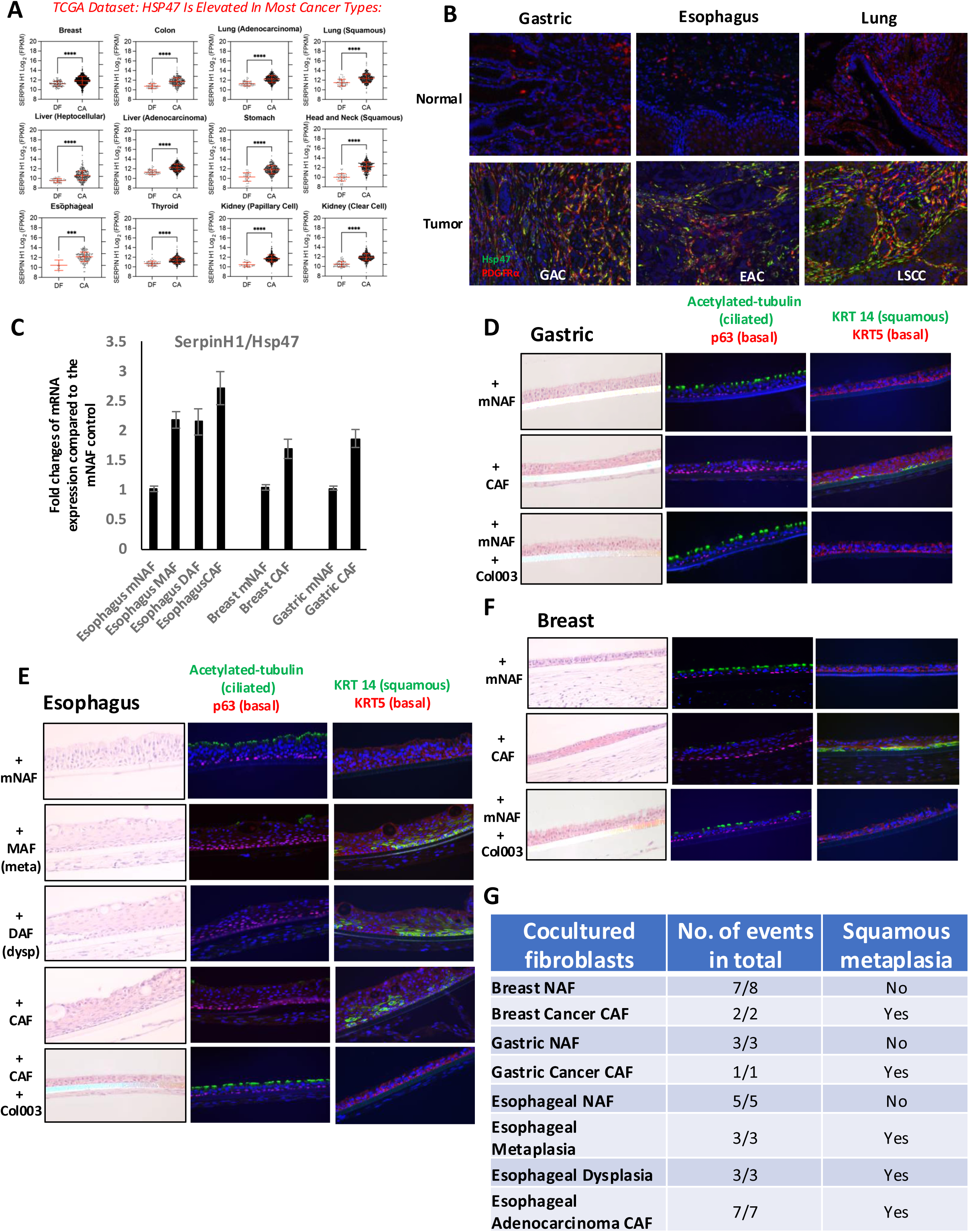
Ability to Induce Metaplasia, Through HSP47 Activity, Develops Early in Progression and is Shared by CAFs from Many Cancer Types. **(A)** Using TCGA data accessed through https://portal.gdc.cancer.gov, we analyzed expression levels of HSP47 (log2[RPKM + 1]) in bulk RNA-sequencing data from 12 cancer types compared to matched histologically-normal controls. Statistical significance was determined using Welch’s t-test. **(B)** Representative immunofluorescence staining images showing Hsp47 and PDGFRα expression in gastric adenocarcinoma (GAC), esophageal adenocarcinoma (EAC), lung squamous cell carcinoma (LSCC) and their adjacent histologically normal tissue specimens. **(C)** The mRNA expression of Hsp47/SERPINH1 was analyzed by qPCR in fibroblasts derived from different cancers or their adjacent histologically-normal or precancerous specimens. **(D-F)** hBECs were cultured in ALI or co-cultured with mNAFs or CAFs derived from gastric adenocarcinoma (GAC) **(D)**; mNAFs, MAFs, DAFs or CAFs derived from esophageal adenocarcinoma (EAC) **(E)**; or mNAFs or CAFs derived from breast cancer **(F)**. Co-cultures were sectioned for histological assessment using H&E staining and immunostained to assess the expression of differentiated epithelial cells and squamous markers. **(G)** Summary of the epithelial phenotypes observed when hBECs were cocultured with fibroblasts from different cancers and disease stages of cancer progression.

### Injury-associated TGF-β Acts Through HSP47-expressing Fibroblasts, Not Epithelial Cells, to Induce Squamous Metaplasia in hBECs

The data above show that ECM remodeling via HSP47 activity is necessary and sufficient for the induction of squamous metaplasia in hBECs, and fibroblasts acquire the potential to induce the creation of precancerous lesions such as squamous metaplasia early in the progression to cancer. Our scRNA-seq analysis revealed a mechanistic clue in that CAFs, which expressed high levels of HSP47, also displayed elevated TGF-β signaling compared to mNAFs, as illustrated by increased expression of TGF-β-targeted ECM proteins and other TGF-β downstream targets including TGFBI, INHBA and periostin (POSTN), whose major activity is to activate TGF-β signaling (Figure S5D). Furthermore, elevated TGF-β signaling in damaged tissue microenvironments upregulates HSP47 expression in fibroblasts ^48–50^. TGF-β signaling is activated by multiple mechanisms, including macrophages and neutrophils responding to inflammatory cues, and by multiple cell types upon injury ^51^ . Since both epithelial cells and fibroblasts are exposed to TGF-β signaling in injured tissue, we asked if signaling in either, or both, compartments was necessary for the induction of metaplasia.

When TGF-β was introduced into the basal medium of the double-sided co-culture system for 21 days, at a time when fully differentiated hBECs were interacting with mNAFs, as we expected, hBECs generated squamous metaplasia (Figure 4A, bottom row, and B). Surprisingly, and in stark contrast, when TGF-β was introduced into the basal medium, at a time when fully differentiated hBECs were cultured alone (in the absence of fibroblasts), hBECs failed to undergo squamous metaplasia (Figure 4A, third row, and B). TGF-b-exposed hBECs exhibited a partial reduction in differentiation markers but no metaplasia. Thus, TGF-β was only able to induce a squamous metaplastic response in hBECs when co-cultured with mNAFs in the double-sided system.

**Figure 4:**
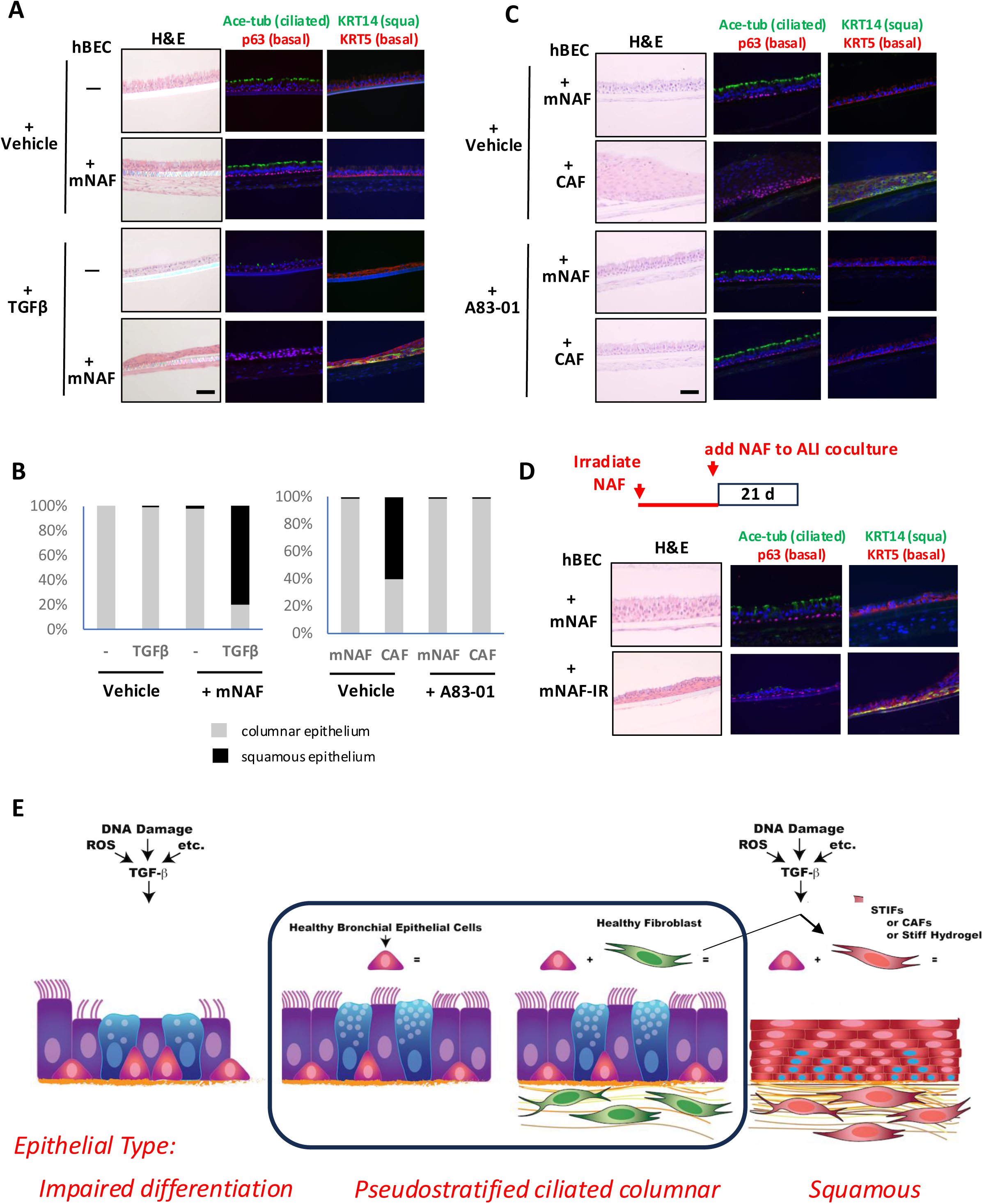
Injury-associated TGF-ß Acts through HSP47-expressing Fibroblasts, not Epithelial Cells, to Induce Squamous Metaplasia in hBECs. **(A)** hBECs were cultured in ALI conditions or co-cultured with fibroblasts under the double-sided condition for 21 days. TGF-β (10 ng/mL) was then added to the medium for 21 additional days. The morphology of the formed epithelium was assessed by H&E staining, and epithelial cell differentiation was assessed by immunostaining for bronchial epithelial and squamous metaplasia markers. **(B)** The percentages of areas in the cultured bronchial epithelium exhibiting squamous metaplasia or pseudostratified epithelium, with or without TGF-β or A83-01 treatment, were quantified and graphed. **(C)** hBECs were co-cultured with mNAFs and CAFs under the double-sided condition in the presence or absence of A83-01 (1 nM) for 21 days. Resulting structures were processed as in **A**. **(D)** Experimental timeline showing fibroblast (mNAFs) irradiation (20 Gy), recovery for 48 hrs in culture, and co-culture with hBECs under a double-sided condition for 21 days (top). Resulting structures were processed as in **A**. **(E)**. TGF-β signaling can act through stromal microenvironment such as instruction of mNAFs (STIFs) or CAFs to create a stiffer stroma that induces squamous metaplasia in hBECs. Conversely, when TGF-β acts directly on hBECs, it represses bronchial epithelial differentiation but does not induce squamous metaplasia.

Cultured mNAFs exposed to TGF-β in 2D culture showed a dramatic upregulation of HSP47 similar to that observed in CAFs. However, TGF-β treatment did not further enhance HSP47 expression in CAFs, which already displayed high HSP47 expression (Figure S7A). Conversely, when TGF-β-exposed mNAFs or cultured CAFs were treated with A83-01, a TGF-β pathway inhibitor, the high expression level of HSP47 was reduced to a similar level as observed in unexposed mNAFs (Figure S7A). To demonstrate the necessity of TGF-β signaling, we treated the double-sided co-cultures with the inhibitor A83-01 and observed that CAFs were no longer able to induce squamous metaplasia (Figures 4B-C and S7B) and instead exhibited a normal pseudostratified structure and ciliated cell differentiation, even in the presence of CAFs. Similar results were obtained using SB431542, another TGF-β pathway inhibitor (Figure S7C-D). These observations further supported that activation of the TGF-β pathway is necessary and sufficient to confer fibroblasts with the ability to induce squamous metaplasia and directly link increased fibroblast HSP47 levels and squamous metaplasia-inducing capability to the activation of TGF-β signaling.

We extended the studies described above by exposing fibroblasts alone to various damage agents before coculturing them with hBECs. Irradiation is known to induce oxidative stress in fibroblasts as well as DNA damage in mammalian tissues^52^. Irradiated mNAFs, just like mNAFs exposed to TGF-β, exhibited a robust upregulation of HSP47 and Collagen I expression at both RNA and protein levels (Figure S7E). When irradiated mNAFs were co-cultured with unirradiated hBECs, squamous metaplasia was induced (Figure 4D). These studies, recapitulated with additional damaging agents (Figure S8), demonstrated that cellular stress associated with chronic inflammation or DNA damage created a fibroblast state exhibiting upregulated HSP47 expression, driving increased collagen fiber content and alignment, elevated tissue stiffness, and subsequent induction of squamous metaplasia in adjacent hBECs. This <u>S</u>tress/<u>T</u>ension-Instructive <u>F</u>ibroblast (STIF) state alone was sufficient to induce hBECs to squamous metaplasia, precursors to tumor formation (Figure 4E).

### CAFs are Sufficient to Induce Squamous Dysplasia When Co-cultured with hBECs Defective in p16 and p53 Pathways

We next explored whether CAFs could further drive progression toward malignancy, i.e could also induce squamous dysplasia. Inactivation of p53 and p16 pathways in epithelial cells, phenotypes commonly observed in early stages of cancers (especially those induced by chronic inflammation and associated with metaplasia and dysplasia), are critical for promoting progression ^53,54^. We therefore abrogated these two tumor-suppressive activities by transducing primary hBECs with HPV E6/E7 (ts^neg^-hBECs) ^55^. Accordingly, the cell cycle inhibitors p21 and p16 were no longer operational in the ts^neg^-hBECs (Figure S9A). Remarkably, the expression of HPV16 E6/E7 proteins did not interfere with bronchial differentiation of hBECs, even after 42 days in ALI culture, whether cultured alone or with mNAFs (Figures 5A and S9C-D) or with trueNFs (Figure S4B). The resulting bronchial epithelial cells (ts^neg^-hBECs) exhibited appropriate epithelial marker expression and low proliferation.

**Figure 5:**
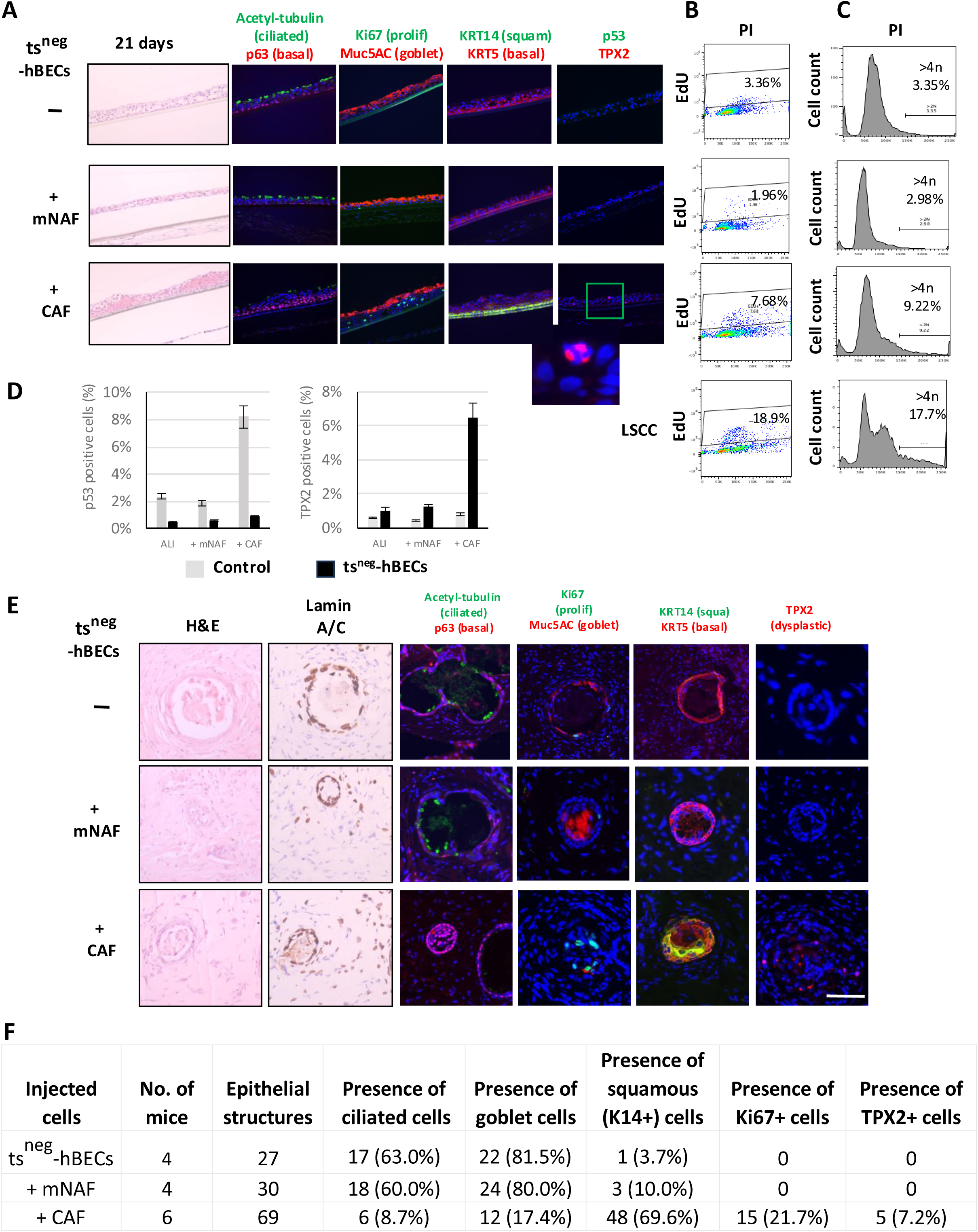
CAFs are Sufficient to Induce Squamous Dysplasia when Cocultured with hBECs Defective in p16 and p53 Pathways both *in vitro* and *in vivo*. (**A**) ts^neg^-hBECs were cultured in ALI or co-cultured with fibroblasts for 21 days. Epithelial structures were stained by H&E and for specific markers: Ki67 (proliferation), Muc5AC (goblet cell), KRT14 (squamous cell), acetylated tubulin (ciliated cell), p63, KRT5 (basal cell), p53 and TPX2, a dysplasia marker. The enlarged image depicts aneuploid cells with three spindle poles identified by TPX2 staining. **(B)** EdU incorporation was measured in ts^neg^-hBECs after co-culture with or without fibroblasts for 21 days and stained with PI and analyzed using flow cytometry. LSCC organoid cells were positive controls. **(C)** The number of cells with chromosome numbers greater than 4N was quantified using PI staining. LSCC organoid cells were positive controls. **(D)** Quantification of p53 and TPX2 positive cells based on immunostaining shown in **(A)**. **(E)** ts^neg^-hBECs were co-injected subcutaneously with different fibroblast populations in immunocompromised NSG mice. After eight weeks, the visualized tissue structures were assessed for morphology by H&E staining and stained for human-specific Lamin A/C to confirm their human origin (left). Differentiation, proliferation, and TPX2 expression were assessed by immunostaining (right). **(F)** Quantification of human epithelial structures and differentiation status based on marker expression profile.

In contrast, when ts^neg^-hBECs were co-cultured with CAFs for 21 days, a squamous morphology was induced, as evidenced by the loss of bronchial differentiation markers, formation of a multilayered esophageal squamous epithelial structure and expression of KRT14 (Figures 5A, S9D-E and S10A). Dramatically, these structures also exhibited phenotypes of dysplasia, including increased proliferation, cellular and nuclear pleiomorphism, aneuploidy, expression of dysplastic markers and disruption of DNA repair responses (Figures 5A-D and S9D-F). Furthermore, after 42 days of co-culture, the 10% of Ki67-positive cells not only resided in the basal layer but also migrated to the apical side, extending through the full thickness of the epithelial layer, closely resembling the pathological and proliferative features associated with human dysplasia *in vivo* including cellular and nuclear pleiomorphism (Figure S9F). Cultures extended to 63 days had robust squamous dysplasia (Figure S9D, bottom right panel). We observed a dramatic increase in EdU-positive proliferating cells, as well as in cells with chromosome numbers exceeding 4N in ts^neg^-hBECs co-cultured with CAFs for 21 days (Figure 5B-C, third panel down from top) compared with cells co-cultured with mNAFs or cultured without fibroblasts (Figure 5B-C, top two panels). As expected, high EdU positivity was noted in LSCC cells cultured in the ALI system (Figure 5B, bottom panel). Notably, the percentage of polyploid ^tsneg-^hBECs co-cultured with CAFs was similar to that observed in LSCC cells (Figure 5C, third panel and bottom panel, respectively). We also investigated expression and localization of TPX2 (Targeting Protein for Xenopus Kinesin-like Protein 2), a spindle assembly factor that plays a crucial role in chromosome segregation during cell division ^56^ and is overexpressed in lung dysplasia and LSCC ^57,58^. In our model, TPX2 expression was significantly increased in ts^neg^-hBECs co-cultured with CAFs but absent in ts^neg^-hBECs co-cultured with mNAFs or cultured alone (Figure 5D, right panel). Notably, some ts^neg^-hBECs co-cultured with CAFs exhibited multiple spindle poles (visible via TPX staining), indicating their aneuploidy (Figure 5A). Taken together, these data further support that ts^neg^-hBECs progressed to dysplasia only in the presence of CAFs, as illustrated by an enrichment in proliferating and aneuploid cells, increased DNA damage responses, morphological changes and increased dysplastic marker TPX2 expression (Figure 5A-D). Importantly, increased TPX2 expression was not observed in hBECs with intact tumor suppressor activity (i.e non-transduced or transduced with a control vector), even when co-cultured with CAFs, despite the induction of squamous metaplasia (Figure 5D).

The observations above were recapitulated *in vivo* when ts^neg^-hBECs were injected subcutaneously alone or combined with different types of fibroblasts and co-injected into mice (Figure 5E). The *in vivo* heterotypic tissue recombinants using ts^neg^-hBECs and CAFs showing the epithelial hallmarks of dysplasia, including increased expression of squamous markers, decreased tissue maturity, uncontrolled proliferation through the full thickness of the epithelial structure, and aneuploidy (Figure 5E-F and S10B). Together, these findings demonstrate that, under these experimental conditions, CAFs are needed to drive dysplastic progression in hBECs even when p53- and p16-dependent pathways are already defective.

### HSP47-Dependent Matrix Stiffness Induces Metaplasia and Controls Progression to Dysplasia by Activating a YAP-dependent Regenerative Program Involving Integrin-mediated Mechanotransduction

Upon tissue damage *in vivo*, the increase in tissue stiffness resulting from deposition of collagen and other ECM proteins, as well as an increase in the alignment of collagen filaments ^46^ has been shown to regulate stem cell activity and plasticity via mechanotransduction ^59,60^, a process by which convergence of physical signals (stiqness, contraction, compression, viscoelasticity, porosity, fiber content, alignment and crosslinking) are converted to biochemical signals to tune transcriptional eqectors of action. To determine if tissue stiffness was a determining factor in inducing squamous metaplasia, we cultured hBECs on matrices of increasing stiffness to mimic potential physiological and pathological conditions. When hBECs were cultured on matrigel (<0.1 kPa) or on Cytosoft matrices with a stiffness of 0.5 kPa, a stiffness corresponding to that of a healthy lung (0.5-2 kPa), expression of bronchial epithelial differentiation genes was high, and expression of squamous and basal cell genes was negligible (Figure 6A). In contrast, when Cytosoft stiffness was increased to levels corresponding to those in diseased lungs (>20 kPa) or higher, the expression of squamous and basal proteins increased significantly, and bronchial epithelial differentiation proteins were markedly reduced (Figure 6A). This demonstrates that matrix stiffness, in absence of any other cell contribution, could activate cell plasticity and induce a squamous metaplastic response in hBECs. Thus, in an injured lung, increased stromal stiffness has the potential to induce squamous gene expression, leading to a squamous metaplastic lesion in the early phase of LSCC progression.

**Figure 6:**
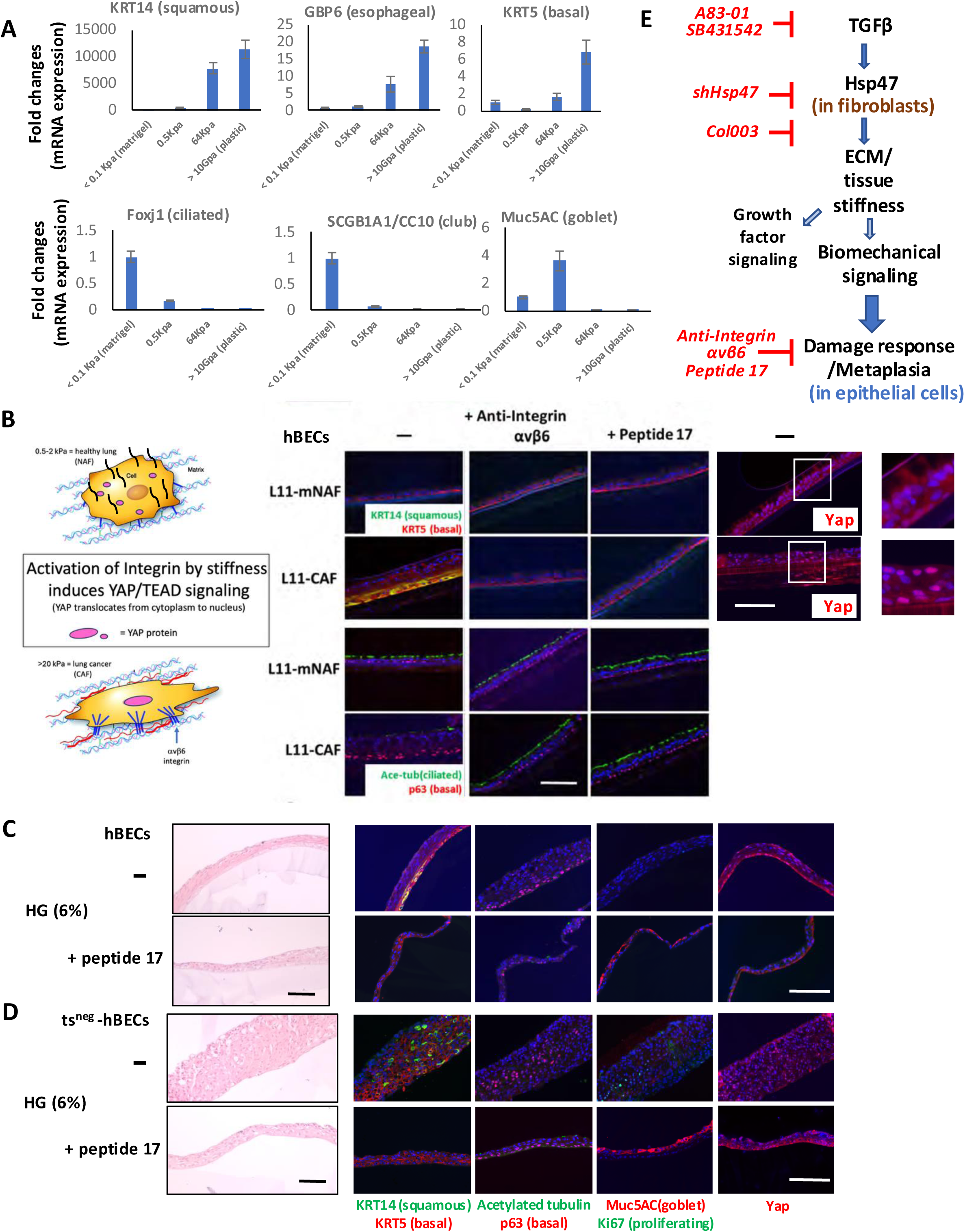
StiLness-induced Activation of Metaplasia is Dependent on Integrin and YAP/TEAD signaling. **(A)** hBECs were cultured on matrices with different stiffnesses (Cytosoft plates). Expression of squamous and bronchial epithelial markers was monitored by qPCR**. (B)** Schematic representation of activation of Yap nuclear translocation in hBECs by stiffer microenvironment (left). hBECs were co-cultured with mNAFs or CAFs in presence or absence of an ITGB6 neutralizing antibody (100 ng/mL) or the YAP/TEAD inhibitor peptide 17 (2 mM) for 21 days. hBECs were assessed for morphology by H&E staining and for differentiation and Yap expression (middle). Magnified insets show the localization of Yap in hBECs (right). **(C)** hBECs were cultured on 6% hydrogel in presence or absence of peptide 17 for 14 days and assessed for morphology by H&E staining (left), and for epithelial cell differentiation, proliferation, and YAP localization status (right). **(D)** The same experimental procedure as in **(C)** was applied to ts^neg^-hBECs.

Integrins mediate complex cell-ECM interactions and are essential for epithelial cells to sense environmental signals and adjust to changes in ECM composition and tissue stiffness ^61^. Because bronchial basal cells are located near the basement membrane and surrounding ECM, we focused our integrin analysis on basal epithelial cells. Examination of RNA sequencing expression profiles of integrin family members in bronchial basal cells identified an elevated class of integrin family members, including integrin αvβ6. Integrin αvβ6 has been recognized as an activator of TGF-β signaling through the release of the active form of TGF-β from its latent complex ^62,63^. Furthermore, mice lacking the integrin β6 (ITGB6) subunit cannot develop pulmonary fibrosis when challenged with the profibrotic agent bleomycin ^63,64^. In our study, blocking integrin αvβ6 with a neutralizing antibody prevented the formation of squamous metaplasia in hBECs co-cultured with CAFs in ALI (Figure 6B). Correspondingly, we observed a dramatic increase in the expression of ITGB6 in the squamous metaplastic regions of the epithelium in ALI (Figure S2A). Notably, expression of integrin β6 was no longer restricted to basal cells, as observed in the normal bronchial epithelium, but was ubiquitously expressed throughout the entire depth of the squamous epithelium. This phenotype was confirmed in clinical metaplastic human lung specimens (Figure S11A). These data demonstrate that integrin αvβ6 could mediate squamous metaplastic responses in hBECs and function as a sensor on epithelial cells in response to a stiffer environment under tissue damage conditions.

The YAP-TEAD pathway is activated in response to a stiffer microenvironment ^65–67^ and plays an important role in regulating squamous metaplastic responses in damaged lungs ^68,69^. In ALI culture, YAP exhibited cytoplasmic localization (i.e., was inactive) in normally differentiated hBECs cultured alone or co-cultured with mNAFs (Figures 6B and S2A). In contrast, when co-cultured with CAFs, hBECs displayed YAP nuclear localization in areas of squamous metaplasia, indicating the activation of YAP (Figures 6B and S2A). Notably, Peptide 17, an inhibitor of the YAP-TEAD pathway, prevented CAF-induced squamous metaplasia (Figure 6B). Moreover, like the HSP47 inhibitor Col003, Peptide 17 could also block TGF-β-induced and irradiation-induced squamous metaplasia in presence of mNAFs (Figure S11B-E). This indicated a critical role for the YAP pathway downstream of HSP47 activity in the regulation of epithelial responses to CAFs or STIFs. Blocking mechanosignaling activated by a stiffer damaged microenvironment efficiently prevented squamous metaplasia in our model (Figure 6E).

To further evaluate ECM stiffness in controling squamous metaplasia, hBECs were cultured on transwells coated with hydrogels at different concentrations (Figure S12A). When grown at low hydrogel concentrations (0.5%), hBECs underwent normal differentiation, forming a monolayer with a pseudostratified ciliated structure and polarized expression of bronchial lineage markers. However, as the hydrogel concentration (and stiffness) increased, hBECs gradually developed metaplasia, that is, they exhibited loss of bronchial lineage marker expression and increases in squamous marker expression (Figure S12A). Notably, ts^neg^-hBECs cultured on high-concentration hydrogel exhibited dysplastic phenotypes, as illustrated by abnormal epithelial morphology, increased proliferation, aneuploidy, and TPX2 expression (Figure S12B). Together, these results demonstrate that heightened tissue stiffness is sufficient for the early progression of LSCC.

Finally, to investigate the mechanistic relevance of mechanosignaling in induction of squamous metaplasia and dysplasia induced by higher concentrations of hydrogel, we assessed activity of the YAP-TEAD peptide 17 inhibitor at a pro-metaplastic hydrogel concentration. YAP activation, as indicated by nuclear localization, was significantly suppressed in the presence of inhibitor, despite the high concentration of the hydrogel (Figure 6C). Administration of inhibitor efficiently prevented expression of squamous markers and restored bronchial epithelial cell differentiation in primary hBECs, even when grown on a stiffer matrix (Figure 6C). Furthermore, a reduction in aberrant proliferation, in addition to restoration of bronchial epithelial differentiation, was observed when inhibitor was added to ts^neg^-hBECs grown on a stiffer matrix (Figure 6D). This further demonstrated that increased matrix stiffness could not only turn on squamous gene expression in adjacent epithelial cells but also directly contributed to the generation of squamous metaplasia and dysplasia through activation of YAP-dependent mechanotransduction. The potential clinical relevance of these observations is high given that current literature documents the poor prognosis that accompanies high HSP47 levels in multiple cancers (Table S2).

### Blocking HSP47 Activities Can Prevent both CAF-induced Squamous Metaplasia and Dysplasia but only Revert Metaplasia

To identify an effective strategy to prevent and/or revert squamous metaplasia and/or dysplasia *in vitro*, we challenged our ALI co-culture model with Col003, an inhibitor that prevents the interaction between HSP47 and collagen^70^. As predicted, the addition of the Col003 inhibitor at the beginning of the co-culture prevented CAF-induced, irradiation-induced or TGF-β-induced squamous metaplasia (Figures 7A and S11B-E). Notably, the establishment of proper pseudostratified bronchial epithelial structures, along with the expression of a ciliated cell differentiation marker, was maintained during Col003 exposure, and induction of KRT14 expression was prevented (Figure 7A). To determine if an HSP47-dependent mechanism was required for maintaining squamous metaplasia once established, we added Col003 after 21 days of co-culture with various fibroblasts (Figure 7B). Following formation of squamous metaplasia, exposure to Col003 for 32 days successfully reverted the squamous metaplasia to bronchial epithelial differentiation in the presence of CAFs (Figure 7B). Notably, exposure to Col003 had no significant effect on the normal differentiation of hBECs cultured alone or co-cultured with mNAFs, even after a longer time in co-culture (Figure 7A-B). Pirfenidone, a clinically approved antifibrotic drug, has been demonstrated to regulate ECM deposition and/or organization by inhibiting HSP47 activity ^71,72^. As expected, pirfenidone had the same effect as Col003 (Figure S13A-B).

**Figure 7:**
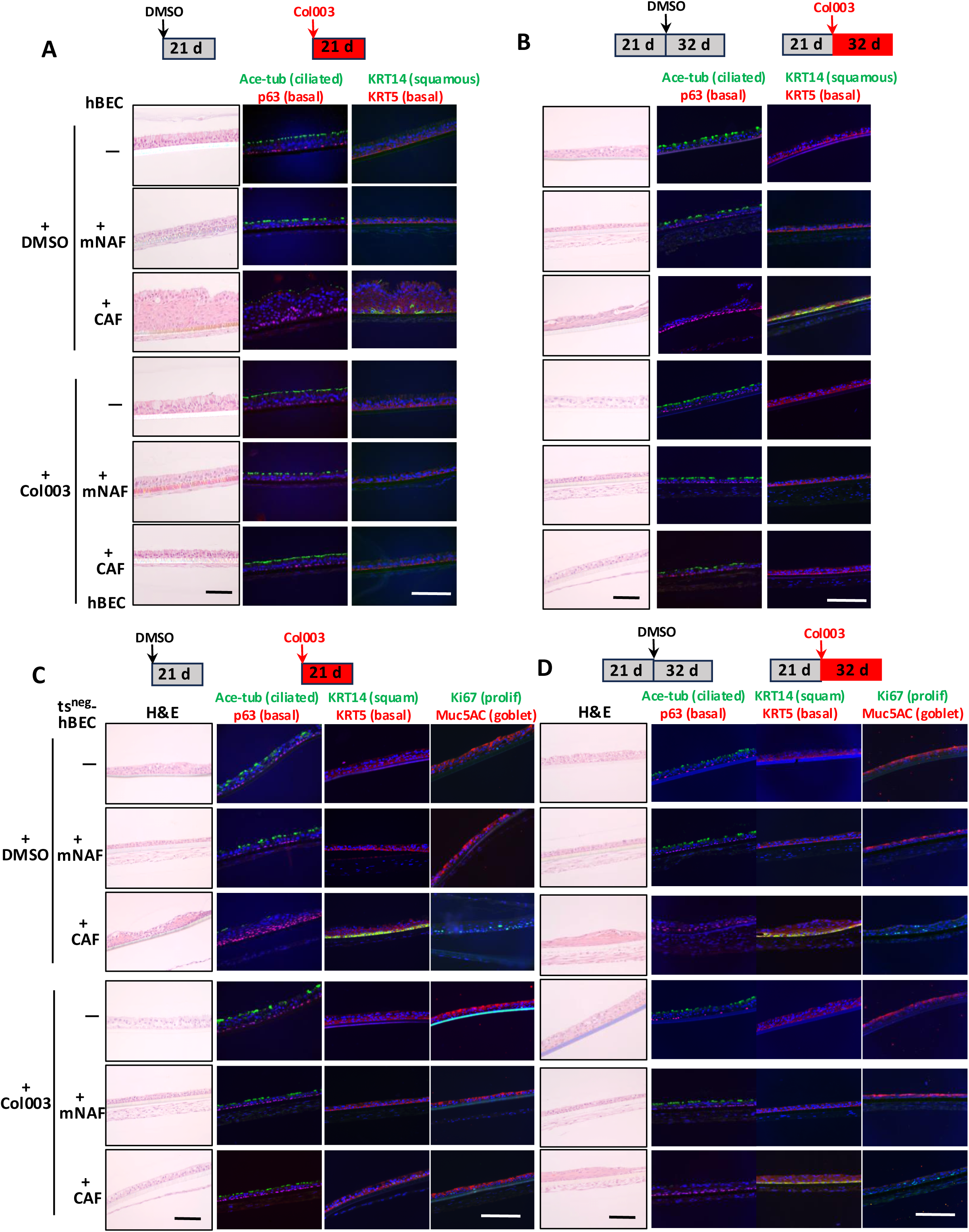
Blocking HSP47-dependent Pathways Effectively Prevents and Reverts Fibroblast-induced Squamous Metaplasia but only Prevents, not Reverts, Fibroblast-induced Dysplasia. **(A)** Experimental timeline showing hBECs co-cultured with mNAFs or CAFs in presence of Col003 (10 μM) or vehicle control for 21 days (top). The formed epithelium was stained with H&E for histological assessment and for expression of squamous and bronchial epithelial markers (bottom). **(B)** Experimental timeline showing hBECs co-cultured with mNAFs or CAFs for 21 days followed by addition of Col003 (10 μM), or vehicle control for 32 additional days (top). The formed epithelium was processed as described in **A** (bottom). **(C)** ts^neg^-hBECs were processed and analyzed as in **(A)**. **(D)** ts^neg^-hBECs were processed and analyzed as in **(B)**.

HSP47-dependent activities, crucial for CAF-induced squamous metaplasia, also exert a critical role in CAF-induced squamous dysplasia, which arises from squamous metaplasia during progression to LSCC. We tested the effects of Col003 at both early and late stages of the ts^neg^-hBEC /fibroblast co-culture. Upon introduction of the inhibitor at the onset of co-culture, efficient prevention of squamous dysplasia was observed in ts^neg^-hBECs (Figure 7C). Indeed, in presence of Col003, ts^neg^-hBECs co-cultured with CAFs underwent normal differentiation, forming a monolayer with polarized expression of ciliated cell markers at the apical side and basal cell markers at the basal side (Figure 7C). Thus, the Col003 HSP47 inhibitor efficiently blocked both pro-metaplastic and pro-dysplastic CAF activity as was also seen with Pirfenidone (Figure S13C). However, the introduction of the Col003 inhibitor after 21 days of co-culture, at a later stage where squamous dysplasia had already been induced in ts^neg^-hBECs by CAFs, failed to reverse the dysplastic process (Figure 7D). These findings suggest that engagement of the HSP47-dependent pathway remained essential for maintaining squamous metaplasia in non-transduced cells, even after its formation, but that it was no longer necessary for sustaining dysplastic phenotypes once dysplasia was established in ts^neg^-hBECs. Pirfenidone showed the same effects as Col003 on CAF-induced dysplasia, prevention at the early stage but failure to revert once established (Figure S13C-D).

## Discussion

### Chronic Stress Induces STIFs, a Tissue-Agnostic Fibroblast State that Choreographs a Regenerative and Onco-enabling Program

Our previous work identified CAFs as a tumor-specific stromal cell population capable of promoting malignant phenotypes in both normal and non-tumorigenic epithelial cells ^18^. Here, we demonstrated that fibroblasts capable of inducing (pre)malignant phenotypes were not only present in the stroma of multiple tumor types but were also present in the stroma associated with pre-malignant lesions (metaplasias, dysplasias and DCIS) and even chronically injured tissues (Figures 1, 3 and S8). Thus, some tumor-promoting signals, previously attributed to CAFs, in fact, develop before tumor formation and may be better described as the acquisition of a STIF state. Elevated HSP47 expression is a hallmark of this state and has also been observed in fibrotic, senescent and aged fibroblasts ^73^ , which are similarly known to promote carcinogenesis ^74,75^.

Multiple insults could induce STIFs, including exposure to physical (IR) and chemical (bile acids) trauma, oxidative stress (tBHP), non-mutagenic carcinogens (talc), genotoxic chemotherapeutic agents (bleomycin) and environmental pollutants (cigarette smoke condensate) (Figures 1 and S8). It is striking to note that the STIF activities are tissue agnostic – they work across tissue types – an important consideration when developing therapeutic interventions. These unanticipated observations highlight that this specific fibroblast state not only plays a critical role in regulating tumor progression and metastasis once tumors are established, as previously reported ^76–82^, but also influences the induction and progression of precancerous lesions during the earliest stages of tumor formation (Figures 1 and 5).

### Mechanistic Insights into the Activation of a Regenerative State and Why this May Lead to the Formation of Pre-malignant Lesions

LSCC represents a group of chronic inflammation-associated cancers (CIACs) that exhibit high rates of metastasis and drug resistance and have poor prognoses. One distinguishing characteristic of early LSCC, or any cancer type associated with chronic inflammation or injury, is the reprogramming of tissue type, generating a metaplastic lesion, and the advancement of malignancy from the reprogrammed lesion (Figure S1A). Why and how this change in developmental state occurs^83,84^ ^85,86^ and how this leads to increased risk for malignant transformation is a compelling question. Understanding metaplasia requires an appreciation of the homeostatic signals that establish and maintain cell identity. During organogenesis, cell identity is directed by an essential “language” involving synergic pathways: classic biochemical gradients of morphogens and growth factors, as well as the more recently discovered instructive cues from the ECM ^87,88^ mechanical signals^60^ and bioelectrical signals^89^. The disruption of tissue homeostasis brought about by injury removes multiple signaling programs that ensure lineage fidelity and stability.

During acute injury, many differentiated cells die, and distinctive basal cell populations temporarily enter an embryonic progenitor state for regeneration of injured tissue. Access to this program is provided by YAP1 translocation to the nucleus where it’s function as a transcription factor activates and represses critical expression networks ^68,69^. Importantly, much of the original mechanical signaling and ECM instruction is still intact and provides the needed guidance to repair, regenerate and restore the original tissue type.

However, during chronic injury or inflammation, key instructive and stabilizing signals become disrupted for an extended time while wound healing is repeatedly attempted. In this context, cells that repeatedly activate YAP1-dependent embryonic programs reside in an environment that lacks not only morphogenic cues, mature cell-cell junctions, and homeostatic biochemical signals but also an extracellular matrix extensively degraded by prolonged protease activity and ongoing stromal remodeling. Concurrently, fibrosis alters the mechanical properties of the stroma. As a result, the subset of cells with an activated regenerative program^90–92^ are forced to proliferate to replace the de-epithelialized area without receiving clear signals for proper reparative organogenesis. Instead, the remaining mechanical cues drive these cells to adopt a phenotype that aligns with the stiffer, altered environment (Figure S14). Mechanical forces have been documented to coordinate these cell fate transitions ^60^. Consequently, in lung squamous cell carcinoma, the loss of ciliated and secretory cells leads YAP-activated basal cells to revert to an embryonic progenitor state that is also observed in the developing bronchial airway and esophagus which derive from the same embryonic progenitor cells in the anterior foregut endoderm ^65–69^. Our work adds compelling and mechanistic evidence that incremental increases in lung stiffness promote the outgrowth of an esophageal tissue state in both the stroma and the epithelium ^42,69^ of a chronically-injured airway (Figures 2 and S4).

Metaplastic squamous cells are better equipped to withstand many of the physical, infectious, and chemical insults that the bronchial airway encounters, often providing protection from further injury and facilitating repair and regeneration within the context of a newly attained metaplastic cell type. However, if the metaplastic change does not fully prevent injury, continuous injury and wound-healing efforts can lead to increased fibrosis and tissue stiffness. When a wound fails to resolve, the tissue remains locked in an embryonic-like progenitor state characterized by persistent wound-healing features that mirror many hallmarks of cancer—such as heightened epithelial proliferation, enhanced survival, increased migration, invasion, dissemination, acquisition of stem cell-like properties, induction of epithelial-to-mesenchymal transition, stiffening of the stromal matrix, localized immunosuppression, metabolic reprogramming, and altered antioxidant gene expression ^93^. Furthermore, dramatic changes in stromal mechanics are transmitted to the nuclear membrane via specific signaling pathways, and excessive nuclear deformation can trigger DNA damage ^94^. Consequently, a mismatch between stromal and epithelial mechanics not only alters epithelial cell identity but also induces genomic damage.

Strikingly, altered mechanical properties of the stroma also activate immune suppression as manifested by naïve T cell proliferation, inhibition of NK and CD8 T cell effector function and induction of suppressive Tregs and MDSCs, which ultimately prevents the immune system from eliminating nascent malignant neoplasms^95–97^. In addition, vascular cells, through mechanosignaling, activate YAP-dependent angiogenesis typically used during wound healing to reconstitute the disrupted vasculature ^98–100^. The convergence of these multiple mechanosignaling programs not only creates a metaplastic lesion but also establishes conditions that explain the increased risk of progression to malignancy. Thus, through mechanosensory cues the STIF state choreographs a multicomponent wound-healing process that, if unresolved, carries the potential for malignancy. Consequently, chronic inflammation contributes far more to cancer development than just the generation of reactive oxygen species and subsequent genetic mutations, as is the current prevailing hypothesis.

### Reconciling Different Perspectives of Cancer Genesis

Early attempts to culture differentiated cells *ex vivo* led to a loss of differentiation markers and functions, even though the cultures retained proliferative capacity ^101,102^. These systems were pivotal in identifying oncogenes, coding sequences that, when activated or overexpressed, can drive cellular transformation and uncontrolled growth. Recent elegant and seminal studies have shown that the expression of many classically identified oncogenes can reprogram primary epithelial cells into tumor precursors by modulating the force transmission between the oncogene-expressing cells and the adjacent ECM ^19,103,104^. For example, mutant Ras signaling increases bundling of actomyosin and causes YAP translocation from the cytoplasm to the nucleus, activating a regenerative program. Remarkably, this effect is conditionally dependent on the tissue microenvironment, since a reduction of ECM-derived force suppresses the manifestation of oncogenic phenotypes in these cells. Thus, the expression of malignant phenotypes driven by expression of oncogenes is conditional and depends on the respective (and combined) force factors at play between epithelial cells and the microenvironmental inputs that modulate them. For decades oncogenes have been studied in cells cultured on plastic surfaces, an experimental condition that resulted in the generation of abnormally high mechanical force (mimicking fibrosis), serving as an unappreciated confounding factor.

Our study demonstrated that increased force transmission from a STIF state can phenocopy oncogene-mediated cell reprogramming that initiates malignancy. Both force factor pathways, those intrinsic to epithelial cells and those extrinsic, that originate from fibroblasts, converge on the YAP-TEAD transcription factor for control of malignant phenotypes. We identified mechanosignaling as a key driver in activating YAP and promoting the initiation and progression of cancers, even in the absence of oncogenic driver mutations in epithelial cells. Strikingly, even when driver mutations are present (ts^neg^-hBECs), mechanosignaling is needed for epithelial cells to exhibit malignant phenotypes (Figure 5). Conversely, we show that increased stromal mechanosignaling, which is extrinsic to the presence or absence of mutations within an epithelial cell, can activate YAP-dependent regenerative programs, thus explaining why not all carcinogens need to be mutagens (Figures 1 and S8). These observations provide a mechanistic basis for previous reports documenting how epithelia bearing oncogenic mutations can only give rise to malignant lesions if placed in an environment with altered biomechanics properties, such as increased tissue stiffness (associated with chronic inflammation), which is associated with increased cell plasticity ^19,104^. These insights, gained from the results of our studies, bring into mechanistic focus years of studies that describe cancer as a disease of dysregulated development ^105^, a wound that cannot heal ^106^, disrupted homeostasis ^107,108^, altered metabolism ^109^, an immune-mediated disease ^110^ and even cancer as a genetic disease ^111^.

### Clinical Relevance

Since STIF signaling can facilitate malignant induction and progression (Figures 1 and 5) and, conversely, stroma engineered to healthy mechanical properties can repress malignant phenotypes (Figures 2 and 7), both aspects documented in the present study, this predicts that standard therapy or precision medicine targeting epithelial cells alone will not be as effective as desired and that re-emergence of disease will be a frequent consequent event. Indeed, YAP-dependent drug resistance is increasingly appreciated^112^. Targeting the stroma, particularly fibroblasts, has long been recognized as a promising strategy to impede tumor growth and progression in numerous experimental models ^113,114^. However, clinical trials have yet to demonstrate the successful implementation of this strategy. In our co-culture model, we convincingly observed that while targeting stromal pathways did not reverse established dysplastic phenotypes, early intervention with this therapeutic strategy effectively prevented dysplasia. These findings underscore the need to reassess the therapeutic efficacy of drugs that have failed at the advanced tumor stage. Our data indicate that the opportunity for prevention exists early in formation of precancerous and cancerous lesions, suggesting potential avenues for clinical breakthroughs.

## Materials and Methods

### Human tissue collection

Fresh tissue specimens, including tumor and histologically normal tissues adjacent to the tumor (matched normal), were prospectively collected from newly diagnosed, treatment-naive patients with LSCC at the McGill University Health Centre. Cancer-free (warm autopsy) healthy lung and esophageal tissue specimens were obtained through the ’Tissue for Research’ initiative, and healthy human esophagus and lung FFPE specimens were acquired through the Cooperative Human Tissue Network Western Division (Nashville, TN) with informed consent under institutional human subject approvals IRB 18-26366 (UCSF) and 2019-5039 (McGill Health Centre). Histological confirmation of tissue diagnosis and staging was independently performed by two expert pathologists, and the samples were labeled accordingly (summarized in Table S3).

### scRNA-seq and analysis

Tumors and matched normal lung tissue specimens from 16 patients diagnosed with LSCC were subjected to scRNA-seq analysis. Our analysis captured transcriptomes of 190,672 individual cells after stringent quality control. We applied the Harmony algorithm to remove batch effects between tissue samples and used the Seurat package in the R programming environment to perform additional preprocessing, dimension reduction, and clustering steps. Unsupervised cluster analysis of the cells produced 10 clusters across two clinical labels: matched normal (103,358 cells) and tumor (87,314 cells). These cells were separated based on gene expression (Figure S1A) into T and NK cells (75,817 cells), myeloid cells (55,591 cells), squamous cells (12,898 cells), mast cells (9,853 cells), alveolar cell types (9,828 cells), ciliated cell types (9,657 cells), fibroblasts (6,115 cells), B cells (5,453 cells), endothelial cells (4,283 cells), and plasma cells (1,177 cells).

### Bronchial organoid derivation

Fresh lung tissue specimens (<40 mg) were minced and dissociated in digestion medium consisting of RPMI 1640 medium supplemented with 10% FBS, 1% penicillin/streptomycin/fungizone (P/S/F)/gentamycin (GA), 0.1 mg/mL DNase I, 0.2 U/mL Liberase. The resulting tissue mixtures were filtered through a 70 mm filter to obtain a homogeneous suspension. The flow-through was centrifuged at 14,000 rpm for 10 min to separate the epithelial component (pellet) from mesenchymal cells (supernatant). The pellets were further digested into single cells using trypsin (0.25%) and passed through a 40 mm filter. Individual cells were collected and suspended in Matrigel (Corning, 356231) at a density of 1x10^6^ cells/50 mL. These cells were then cultured in bronchial organoid medium for 14-21 days ^32^. The resulting organoids were passaged and expanded for further analysis. To stain cells grown as cultured organoids, they were fixed in 4% paraformaldehyde for 1h at room temperature (RT). After fixation, organoids were collected and embedded in histogel for further processing following a standard FFPE embedding procedure.

### Primary fibroblast culture

We have extensively characterized the conditions that permit human endothelial, pericyte, adipocyte, and fibroblast growth in culture in several publications, starting with our original recombinant co-culture studies in 1999 ^18^. The initial enrichment for fibroblasts occurs when we centrifuge a single-cell suspension at 16,000 RPMs for 10 minutes. Adipocytes, immune and epithelial cells do not pellet. The cell pellet was then washed once in FGM2 medium (Lonza, CC3132) and cells were seeded in one well of a six-well plate for further expansion in FGM2 medium. The culture conditions used to expand the fibroblasts for the recombinant experiments do not support the survival or growth of endothelial cells or pericytes (they need VEGF). Figure S3B, provides data characterizing the fibroblasts that were used and show a lack of staining for pericyte and endothelial markers and expression of PDGFRα, a fibroblast marker.

### Bronchial epithelial cell ALI (air-liquid interface) culture

Bronchial organoids grown in Matrigel were harvested and digested using TrpLE to remove Matrigel and dissociated into single cells. 10^5^ cells were seeded in a 6.5 mm transwell insert (Corning, 3470) and cultured for 2-4 days in DMEM/F12 (3.5:1.1), 10% FBS, 0.18 mM adenine, 0.5 µg/mL hydrocortisone, 5 µg/mL insulin, 10^-10^ M cholera toxin (Sigma, C8052), 10 ng/mL EGF, and 1X Primocin until a confluent layer was formed. The growth medium was then removed from the inner and outer chambers of the transwell. The outer chamber was supplied with differentiation medium (Stem Cell Technologies Inc., 05001), while the inner chamber was exposed to air for 21 days to allow spontaneous differentiation of bronchial epithelial cells. The differentiation process was monitored by measuring the TEER values using a Millicell ERS-2 Voltohmmeter.

For studies involving damage-associated factors or inhibitor treatment, differentiation medium was supplemented with specific factors during the differentiation process. Alternatively, bronchial epithelial cells were fully differentiated after 21 days of ALI culture and then exposed to the factors for the indicated time periods. The formed bronchial epithelium was directly collected for RNA analysis or fixed in 4% paraformaldehyde for 1h at RT for histological analysis or immunofluorescence staining.

### Epithelial cell /fibroblast ALI co-culture

For the single-sided co-culture, 2x10^5^ fibroblasts were seeded onto the fibronectin-coated bottom of the outer chamber one day before adding epithelial cells to the inner chamber. Bronchial epithelial cells were cultured in bronchial epithelial growth medium for two to four days until they reached confluence. Subsequently, ALI culture was initiated by switching to bronchial differentiation medium. The tissue recombinant was then fixed in 4% paraformaldehyde for one hour at RT for histological analysis or immunofluorescence staining.

For the double-sided co-culture, the outer chamber of the transwell membrane was coated with fibronectin (10 mg/mL) overnight at RT. The transwell insert was then inverted and placed into a P150 culture plate. 2x10^5^ fibroblasts were seeded on the top surface of the inverted transwell insert and allowed to attach to the membrane at 37°C for four hours. Subsequently, the transwell insert with the attached fibroblasts was transferred back to the original 24-well plate. Fibroblasts were cultured overnight after the addition of FGM2 medium in both outer and inner chambers. Once the fibroblasts were established, bronchial epithelial cells were added to the inner (top) chamber. The medium in both chambers was then switched to bronchial growth medium for two to four days until the epithelial layer reached confluence. Subsequently, the ALI co-culture was initiated by switching to the differentiation medium. The resulting tissue recombinant was fixed in 4% paraformaldehyde for one hour at RT for further histological analysis or immunofluorescence staining. For single-sided co-culture, fibroblasts were seeded at the bottom of the outer chamber and cultured overnight. Epithelial cells were seeded into the inner chamber on the second day.

### Generation of tissue recombinants in mice

2.5 x10^5^ bronchial epithelial cells derived from bronchial organoids in combination with or without 5x10^5^ fibroblasts were resuspended in 50 mL of 50% Matrigel. The cell mixture was then injected subcutaneously into the right hind legs of nine-week-old female NSG mice. The engrafted tissues were collected four or eight weeks after cell injection for histological and immunohistochemical analyses. Mice injected with 5x10^5^ fibroblasts without epithelial cells were used as controls. Experiments were carried out under institutionally approved animal protocol AN-177694/192445.

### LSCC organoid culture

LSCC organoids (LTKL) were cultured in a 3D Matrigel matrix and supplied with DMEM/F12 medium supplemented with 20 ng/mL EGF, 1x N2 supplement, 1x B27 supplement, 10 μM Y27632, 50 ng/mL FGF2, and 1x Normocin.

### Matrisome analysis using a proteomic approach

Tumor and matched normal lung tissue specimens from 21 patients diagnosed with LSCC were subjected to ECM extraction and proteomic analysis as previously described ^43^. Fold changes in tumor versus matched normal tissues for individual patients were calculated.

### Decellularization of Fibroblasts

Fibroblasts (2 × 10⁵) were seeded in the inner chamber of a 24-well transwell and cultured for 7 days. Decellularization was performed by treating the cells with 300 µL of 20 mM NH₄OH and 0.5% Triton X-100 in PBS for 5 minutes. The solution was then removed, and the decellularized ECM was washed twice with PBS. To eliminate residual nucleic acids, the ECM was treated with 200 µg/mL DNase and RNase for 30 minutes at room temperature. The ECM was then washed three times with PBS before seeding 1 × 10⁵ hBECs for culture under ALI conditions.

### Histology and immunofluorescence staining

The tissue structures formed from the ALI culture were sectioned longitudinally, whereas the other tissues were sectioned using the regular method. Tissue sections were then subjected to histological staining using a standard H&E protocol or immunofluorescence staining by incubating the tissue sections with specific primary antibodies and, subsequently, fluorescence-conjugated secondary antibodies. The primary antibodies and dilutions used for immunofluorescence staining were as follows: acetylated Tubulin (Santa Cruz Biotechnologies (SCBT), sc23950, 1:100), p63 (Cell Signaling, 13109, 1:300) CC10 (SCBT, sc365992, 1:100), KRT5 (Abcam, ab64081, 1:500), Muc5AC (Abcam, ab198294, 1:250), Ki67 (Dako, M7240, 1:200), Involucrin (Abcam, ab68, 1:100), YAP (Cell Signaling, 140743, 1:200), Integrin β6 (Cell Signaling, 9513, 1:200) KRT14 (Biocare, CM185C, 1:100), Collagen I (Abcam, 1384921, 1:1500), p53 (Cell signaling, 488183, 1:100) SPC (Abcam, ab90716, 1:100), Collage IV (Abcam, 6586, 1:400), ZO-1 (Invitrogen, 33-9100 1:500), HSP47 (Abcam, 109117, 1:400), GPB6 (Sigma, HPA027744, 1:500), KRT10 (Abcam, ab9026, 1:100), PDGFRα (Cell signaling, 3164 1:100), alpha-Smooth Muscle Actin (Dako, 0851, 1:400), 53BP1(Cell Signaling, 88439, 1:200), p21 (Cell Signaling, 2947, 1:200), (TPX2 (Sigma, HPA005487, 1:200) and HPV16 E6 antigen (SCBT, SC460, 1:100)

### Multiplex immunohistochemistry (mIHC)

Immunostaining of 5-micron thick formalin-fixed paraffin-embedded human tissue specimens (summarized in Table S3) was carried out using the Tyramide Signal Amplification (TSA) technology. M*odule 1*: HSP47 (primary antibody dil. 1/2,000, 1 hr, RT) was detected with goat anti-rabbit MACH2 (30 min, RT, Biocare Medical), FITC dye (2 min, RT, Akoya Biosciences), and KRT14 (primary antibody dil. 1/500, overnight, 4 °C) was detected using anti-mouse MACH2 (30 min, RT) and Cy3 dye (3 min, RT); anti-collagen 1 (dil. 1/5,000, 1 hr, RT) was detected using goat anti-rabbit MACH2 (30 min, RT) and Cy5 dye (10 min, RT). *Module 2*: anti-acetylated tubulin (dil. 1/500, 1 hr, RT) detected with goat anti-mouse MACH2 (30 min, RT) and FITC dye (2 min, RT); anti-KRT14, as described for module 1; anti-GBP6 (dil. 1/750, 1 hr, RT) was detected using goat anti-rabbit MACH2 (30 min, RT) and Cy5 dye (10 min, RT). M*odule 3*: Integrin β6 (dil. 1/1000, overnight, 4 °C) and detected with goat anti-rabbit MACH2 (30 min, RT, Biocare Medical) and Cy3 dye (3 min, RT). KRT14 (primary antibody dil. 1/500, 1 hr, RT) was detected using anti-mouse MACH2 (30 min, RT) and Cy5 dye (10 min, RT).

### Imaging

H&E-stained slides were imaged at 20x magnification using an Aperio AT2 whole-slide scanner (Leica). Slides stained for mIHC modules were imaged at a high resolution at 20x magnification (tissue mIHC) or 40x (ALI immunofluorescence) using a BZ-X800 fluorescence microscope (Keyence). At least five independent 20x fields were typically acquired for each specimen. A scale bar of 100 um is labeled in each set of images.

### Quantitative Analyses

*<u>Immunofluorescence:</u>* The percentages of epithelial cells of different lineages were quantified by positive cell counting based on marker expression. Collagen filament coherence was quantified using OrientationJ based on collagen staining . *mIHC:* Images including cytokeratin 14-positive lesions were processed as 8-bit merged image files and then as tiff composite image files using Image J/Fiji (version 2.1.0/1.53c). Composite images were then analyzed using the QuPath software (version 0.2.3). Briefly, stromal areas were annotated and queried at full resolution using a Gaussian prefilter for positive pixels for FITC (HSP47) and Cy5 (collagen 1) using fluorescence intensity thresholds (HSP47:20; collagen 1:35) defined after the analysis of control true normal lung specimens (Figure S15).

### Immunoblotting and Real-time qPCR

Immunoblotting was performed as previously described . Total RNA was isolated from cells and reverse-transcribed into cDNA as previously described . qPCR (Taqman) was performed in triplicates using the delta-delta CT method with gene-specific primer/probe sets. The expression of the glucuronidase B (GUSB) housekeeping gene was used to normalize the variance in the input cDNA.

### Retroviral gene transfer

Lentiviral plasmids expressing Myc-DKK tagged HSP47/SERPINH1 (OriGene: RC209847L3) or short hairpins (OriGene: TL316478B) for HSP47/SERPINH1 and matched control plasmids were used to transfect packaging 293 cells. Lentiviral suspensions were used to transduce various fibroblast populations for 48 hours. Transduced cells were selected for 48 hours with 2 mg/mL puromycin and collected for Western blot analysis or used for the double-sided ALI co-culture.

### Epithelial cell culture on matrices with different stiffnesses

For testing of hBECs on Cytosoft plates (Advanced Biomatrix) with different stiffnesses, bronchial organoids were dissociated into single cells. 3x10^5^ bronchial epithelial cells were seeded and cultured for seven days in bronchial epithelial growth medium. Cells were then collected for further analysis.

For testing of hBECs on different concentrations of hydrogel, bronchial organoids were dissociated into single cells. 2x10^5^ bronchial epithelial cells were seeded at different concentrations of hydrogel prepared using the HyStem-C hydrogel kit (Advanced Biomatrix). In brief, Extralink, Glycosil and Gelin-S were added in a 1:2:2 volume ratio. Dilutions of each component were adjusted to obtain concentrations of hydrogels corresponding to different stiffnesses ^19^.

### Statistical Methods

A two-sided t-test assuming unequal variance was used to assess the relationships between mRNA expression levels of Serpin H1/Hsp47, Collagen I, KRT14, GBP6, KRT5, Foxj1, SCGB1A1, and Muc5AC; protein expression of Hsp47; cell counts in immunofluorescent images; and TEER measurements. A threshold of P ≤ 0.05 was considered statistically significant. Data are presented as mean ± SEM.

## Acknowledgments

We thank Dr. Calvin Kuo (Stanford University, California) for generously sharing a lung squamous cell carcinoma organoid culture and Dr. Jeroen Roose, (UCSF) for advice on bronchial organoid culture. We thank Drs. D. Jackson, L. Norton, L. Van’t Veer, A. Ashworth and G. Lewis for support and Dr. B. Emerson for critical reading of the manuscript. We acknowledge the CRUK STORMing Cancer team for helpful advice and discussions. We dedicate this work to Zena Werb, Mina Bissell and Curtis Harris pioneers studying stroma and chronic inflammation.

## Author contributions

D.P., P.G., S.H.., B.S., L. F. and T.D.T. designed research; D.P., P.G., C. T., J.A.C., V.S., N.B, S.C., J.B., S.O., M.K.S., D.L.G., J.B., J.B.B., J.P. R., S.S., R.B., S.L., D.R.F., A.U. and I.R. performed research; D.P., P.G., J.A.C. and T.D.T. analyzed data; and D.P., P.G. and T.D.T. wrote the paper.

## Data Availability Statement

Raw data and complete MS data sets have been uploaded to the MassIVE repository of the Center for Computational Mass Spectrometry at UCSD and can be downloaded using: ftp://MSV000095825@massive.ucsd.edu or via the MassIVE website: https://massive.ucsd.edu/ProteoSAFe/private-dataset.jsp?task=719da92aa46e44659273fe044f335dfb (MassIVE ID: MSV000095825; ProteomeXchange ID: PXD055769). [for MS data access in MassIVE (UCSD): Username: MSV000095825_reviewer; Password: winter].

Single-cell RNA-seq data and de-identified patient metadata have been deposited on Figshare: https://doi.org/10.6084/m9.figshare.26997133.

---

**Supplementary Figure S1:**
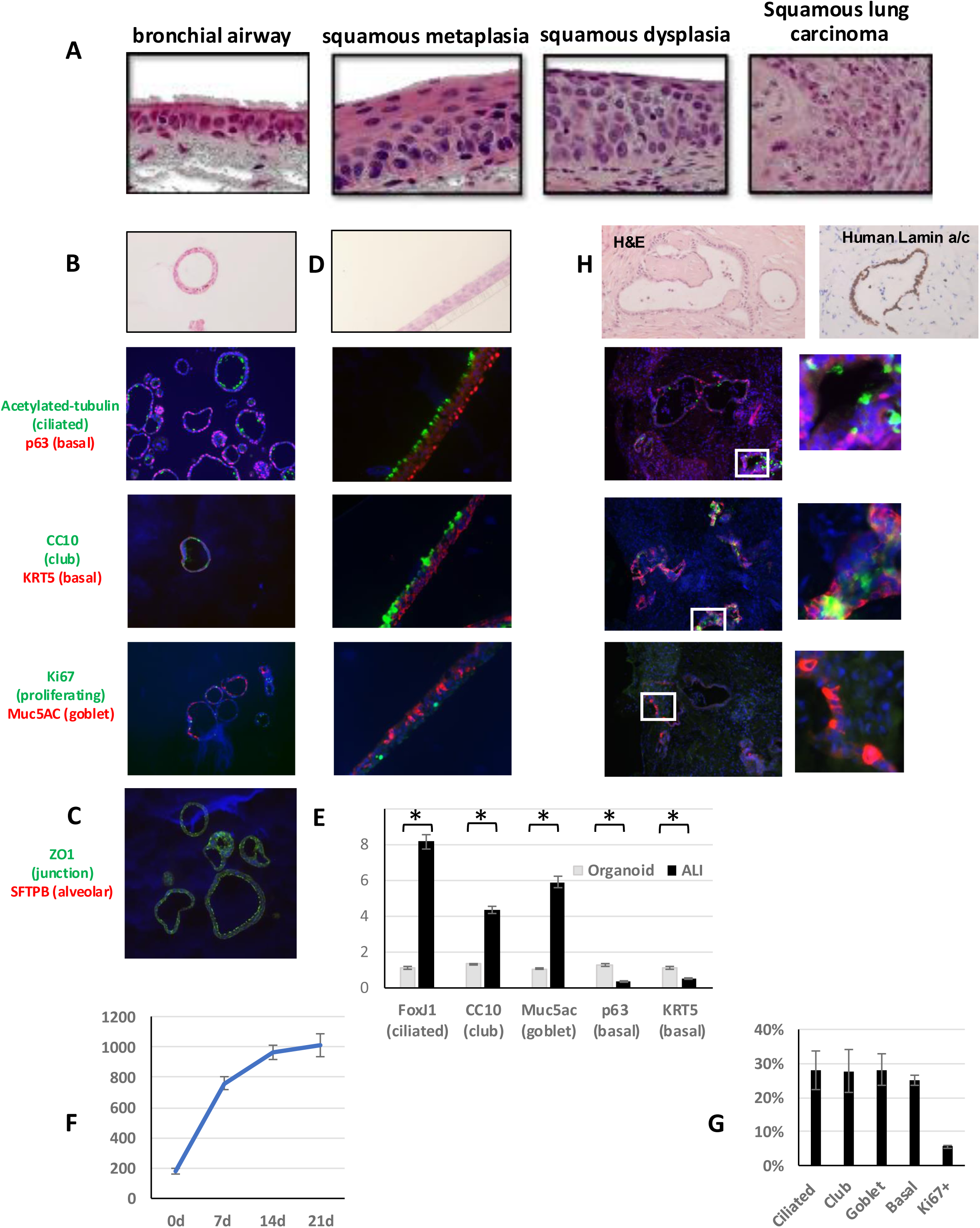
Recapitulating Representative Fully DiLerentiated and Functional Human Bronchial Airway Epithelial Phenotypes In Vitro in Air Liquid Interface and In Vivo. **(A)** Representative H&E stainings of human specimens showing normal bronchial epithelium, squamous metaplasia, squamous dysplasia and LSCC. **(B)** hBECs derived from bronchial organoids were cultured in Matrigel or **(D)** cultured in the ALI (air-liquid interface) condition for 21 days. The morphology of the formed epithelium was assessed by H&E staining and differentiation status was assessed by immunostaining of bronchial epithelial markers (acetylated tubulin for ciliated cells, CC10 for club cells, Muc5AC for goblet cells, p63 and KRT5 for basal cells, Ki67 for proliferating cells). **(C)** Cultured bronchial organoids were immunostained for ZO-1, a tight junction protein, and SFTPB, an alveolar cell marker**. (E)** mRNA expression of differentiation markers was compared by qPCR in hBECs cultured as organoids or as ALI. **(F)** Epithelial permeability was assessed using TEER measurements during airway epithelial differentiation under ALI conditions. **(G)** After culture in ALI, the percentage of each cell lineage was quantified based on immunostaining.**H )** hBECs derived from organoids were subcutaneously injected into immunocompromised NSG mice. After four weeks, tissue structures were assessed for morphology by H&E staining and for confirmation of human origin by immunostaining with a human-specific Lamin A/C antibody (top). Epithelial cell differentiation was assessed by immunostaining for bronchial epithelial markers (bottom)

**Supplementary Figure S2:**
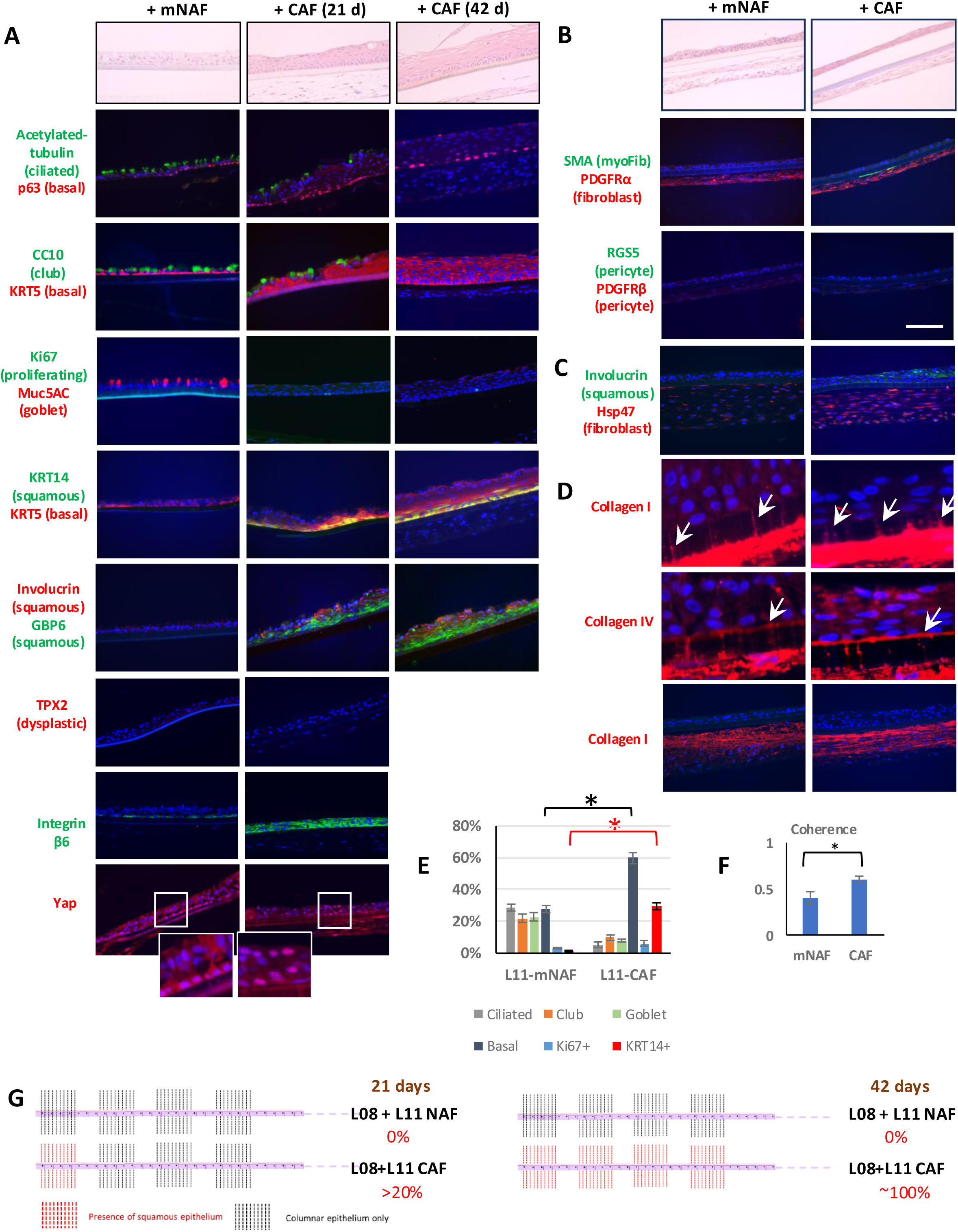
Representative Phenotypes of Fully Differentiated, Functional Human Bronchial Airway Structures and Squamous Metaplasia In ALI Tissue Recombinants *In Vitro*. For this summary, examples of staining were obtained from multiple co-cultures. **(A)** hBECs derived from organoids were co cultured in ALI as homotypic recombinants (both epithelial cells and fibroblasts were obtained from the same tissue state (histologically-normal) or as heterotypic recombinants (epithelial cells were obtained from histologically-normal tissue and fibroblasts from progressive stages of cancer) for 21 or 42 days. The morphology of the formed epithelium was assessed by H&E staining and differentiation status by immunostaining for bronchial epithelial markers. Induction of squamous metaplasia was assessed by H&E staining and differentiation was assessed by immunostaining for squamous epithelial markers (Involucrin and KRT14). Esophageal embryologic origin of lung squamous metaplasia reported *in vivo* ^37,38^ was confirmed by immunostaining of *in vitro* cocultures for the esophagus-specific marker GBP6. Markers for dysplasia (TPX2 and aneuploidy) and for mechanotransdcution pathway (Integrin b6 and nuclear translocation of YAP) were also assessed by immunostaining in cocultures. **(B)** hBECs were co-cultured with fibroblasts obtained from histologically normal tissue adjacent to cancer (matched Normal-Associated Fibroblasts; mNAFs) or from cancer lesions (Carcinoma-Associated Fibroblasts; CAFs) for 21 days. The morphology of the formed epithelium was assessed by H&E staining (top). Cocultured fibroblasts were assessed for expression of fibroblast markers; PDGFRα was expressed in both mNAFs and CAFs, while both lacked the expression of the myofibroblast-specific marker alpha-smooth muscle actin (SMA) and pericyte markers RGS5 and PDGFRb **(C)** The expression of the squamous epithelial marker (Involucrin) in hBECs and of the fibroblast marker (Hsp47) was assessed by immunostaining in cocultures. **(D)** The fibroblast-generated ECM network was assessed by collagen staining. Interactions of hBECs with fibroblasts through ECM proteins in double-sided co-culture were visualized using overexposed Collagen I or Collagen IV staining, with arrows indicating collagen fibrils crossing the transwell pores and interacting with epithelial cells. **(E)** The percentage of each cell lineage in cocultures was quantified based on immunofluorescence staining. **(F)**The coherence of collagen filament alignment was quantified in collagen stained specimens in **(B)** using OrientationJ in ImageJ. **(G)** Schematic images showing serial sections across the entire Transwell membranes (6.5 mm diameter) after 21 days (left) or 42 days of co-culture (right). Vertical dashed lines represent the sections used for H&E staining. The presence of squamous metaplasia is indicated by red dashed lines. The overall percentage of surface areas with evidence of squamous tissue is shown.

**Supplementary Figure S3:**
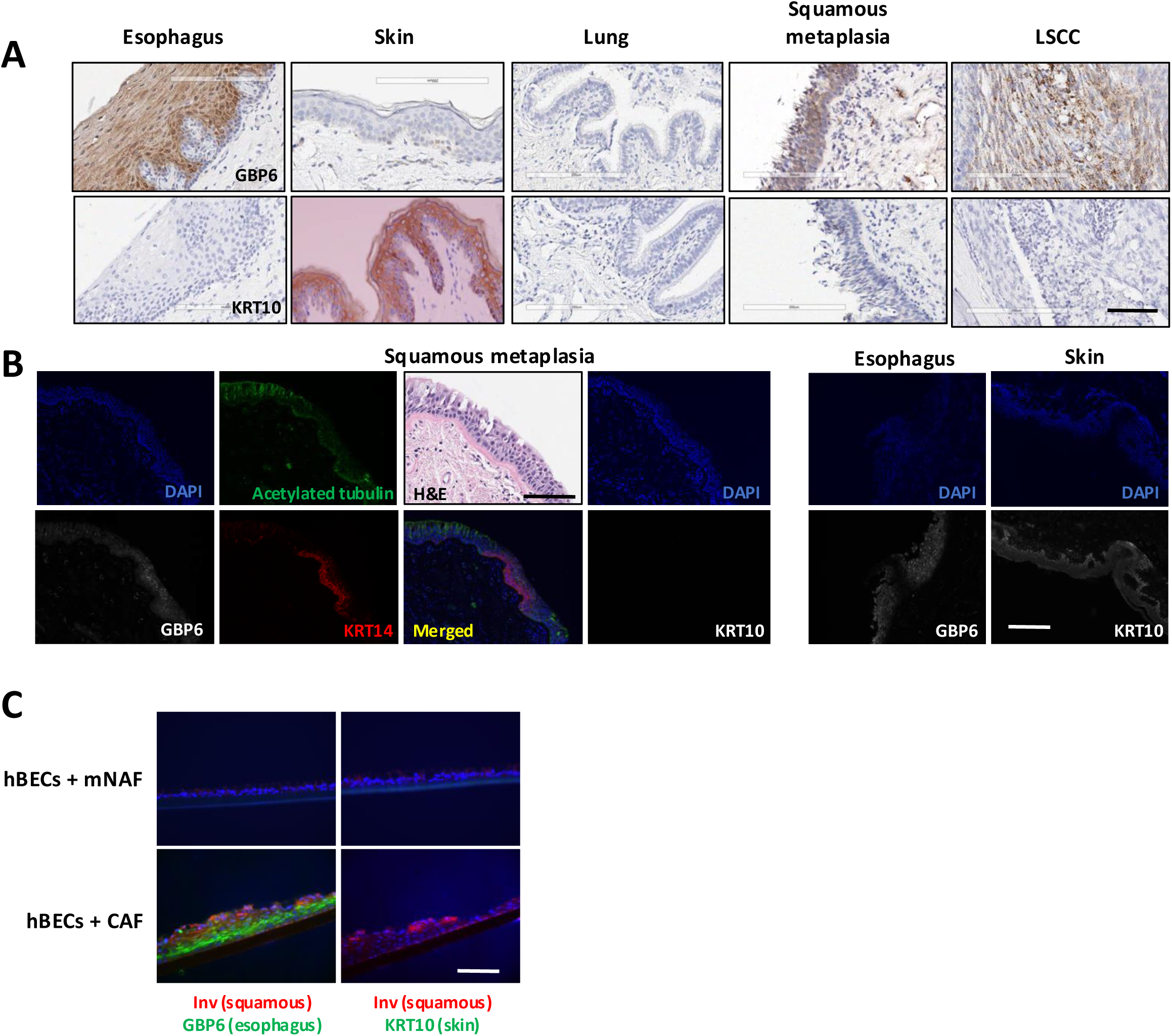
Lung Squamous Metaplasia Exhibits Features of Esophageal Squamous Epithelium, which Shares a Common Embryonic Progenitor with Lung Tissue. **(A)** Immunostaining of human disease-free esophagus, skin lung, lung squamous metaplasia and LSCC specimens for esophagus-specific marker GBP6 or skin-specific marker cytokeratin 10 (KRT10) using DAB (brown). Disease-free lung does not express either of the two markers. Lung squamous metaplasia and LSCC both express the esophageal marker GBP6 but not the skin marker KRT10. **(B)** Immunofluorescence-based staining of lung squamous metaplasia specimens from cancer-free patients for ciliated cell (acetylated tubulin), squamous cell (KRT14), esophageal (GBP6) and skin (KRT10) markers (left panel). The esophagus or skin was stained as a positive control (right). **(C-D)** immunostaining-based characterization of esophageal-like squamous metaplasia in hBECs **(C)** or ts^neg^-hBECs **(D)** cocultured with mNAFs or CAFs.

**Supplementary Figure S4:**
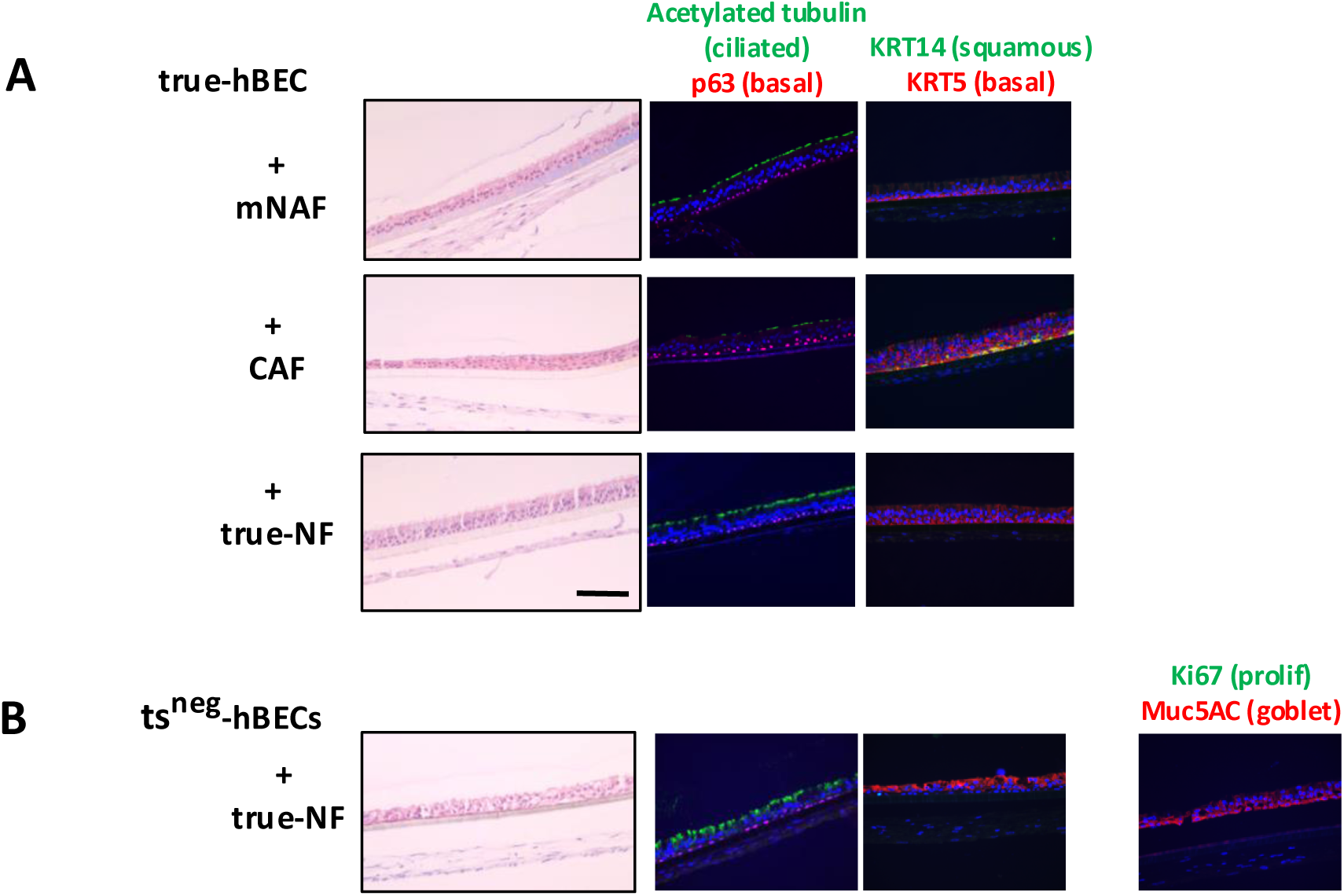
Fibroblasts and Epithelial Cells from Individuals that Have Not Had Cancer (true-NF and true-hBECs) Recapitulate Phenotypes Obtained with mNAFs and hBECs. **(A)** hBECs derived from disease-free lungs (true hBEC) were co-cultured with mNAFs, CAFs, or tNFs (true normal fibroblasts derived from disease-free lungs). The morphology of the formed epithelium was assessed by H&E staining, and epithelial cell differentiation status by immunostaining of bronchial epithelial and squamous markers. **(B)** ts**^neg^-**hBECs were cocultured with tNFs and processed as in **A**. The proliferation of epithelial cells was assessed by immunostaining of Ki67.

**Supplementary Figure S5:**
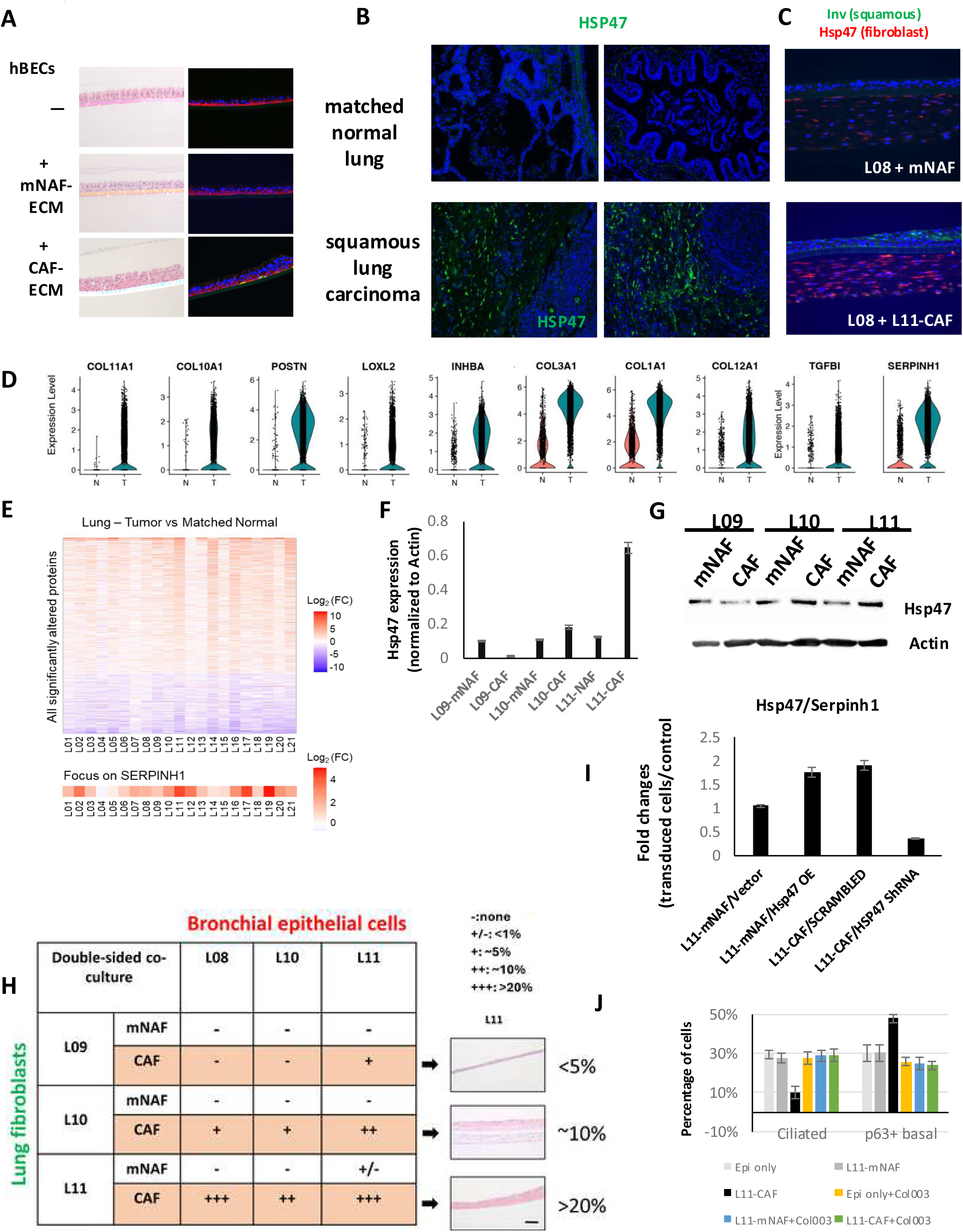
ECM Remodeling is Necessary and Sufficient for Induction of Squamous Metaplasia in hBECs. **(A)**. hBECs were cultured on decellularized ECM derived from mNAFs or CAFs. The morphology of the formed epithelium was assessed by H&E staining, and epithelial cell differentiation by immunostaining of bronchial epithelial and squamous markers. **(B)** Representative images for immunofluorescent staining for Hsp47 in LSCC and their adjacent histologically normal lung specimens. **(C)** hBECs were cocultured with mNAFs or CAFs for 21 days. The expression of the squamous marker (Involucrin) and the fibroblast marker (Hsp47) was assessed by immunofluorescence staining. **(D)** Violin plots showing the expression of the most differentially expressed genes in mNAFs versus CAFs, based on scRNA-seq analysis. **(E)** Differentially expressed matrisomal proteins in LSCC compared to matched normal lung specimens were identified by proteomic mass spectrometry in 21 LSCC patients. (q-value ≤ 0.01 and |log2(FC)| ≥ 0.58). A heatmap for SERPINH1 is displayed (bottom; q-value = 3.03 e-51). **(F)** Quantification of HSP47/SERPINH1 expression levels in cultured CAFs and mNAFs from different LSCC patients by western blotting. Actin was used as a loading control for normalization. **(G)** The images of western blotting were shown. **(H)** Summary of epithelial-fibroblast co-cultures from multiple LSCC patients, showing varying sensitivities to squamous metaplasia induction. Representative images of H&E staining for different levels of squamous metaplasia are shown on the right. **(I)** The mRNA expression for Serpin H1/Hsp47 was compared by qPCR in fibroblasts transduced with different lentivirus constructs. **(J)** Quantification of cell lineage representation in cocultures in presence or absence of 10 μM of Col003 based on immunostaining.

**Supplementary Figure S6:**
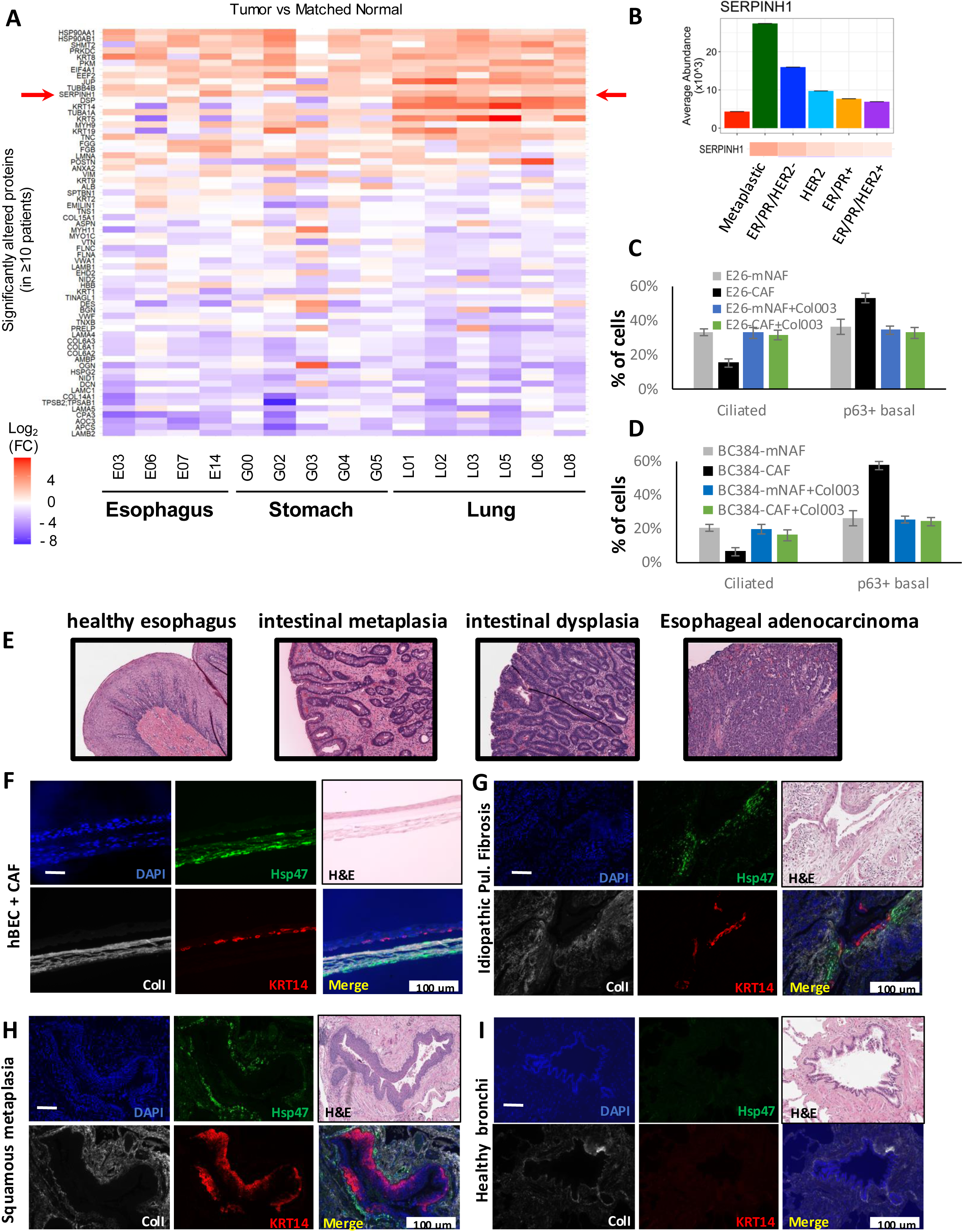
Elevated Expression of HSP47 Detected in Malignant and Pre-malignant Tissues. **(A)** A group of consistently altered matrisomal proteins was identified by proteomic mass spectrometry when comparing cancer to matched histologically normal tissue specimens in multiple patient specimens from three CIAC cancer types (q-value ≤ 0.01 and |log2(FC)| ≥ 0.58). Serpin H1/Hsp47 was consistently upregulated in the cancer matrisome across all three cancer types. Heatmap showing the log_2_-fold protein changes (‘Tumor’ vs. ‘Matched Normal’) for each individual patient assessing the proteins that were significantly altered in at least 10 patients (q-value ≤ 0.01 and |log_2_(FC)| ≥ 0.58). The arrow points to SERPINH1/Hsp47. **(B)** The abundance of Serpin H1/Hsp47 specific peptide detected in ECM derived from different breast cancer types were plotted based on ECM analysis. **(C)** Quantification of cell lineage representation in hBECs cocultured with mNAFs or CAFs derived from EAC, in the presence or absence of 10 μM of Col003, based on immunofluorescence staining. **(D)** Quantification of cell lineage representation in hBECs cocultured with mNAFs or CAFs derived from ER-positive breast cancer, in the presence or absence of 10 μM of Col003, based on immunofluorescence staining. **(E)** Representative H&E stainings of human specimens showing progression from healthy squamous esophagus to intestinal metaplasia, dysplasia and esophageal adenocarcinoma**. (F)** hBECs co-cultures with CAFs using the double-sided system were stained for KRT14, HSP47, and Collagen I using multiplex IHC. **(G-H)** Immunohistochemistry of lung tissues at higher risk for developing lung cancer mimics the colocalization of markers seen in the ALI co-cultures with an exhibition of focal areas high in HSP47 adjacent to increased collagen content and squamous metaplasia (expression of KRT14) in the nearby epithelium. Expression of these markers was not observed in healthy human bronchial tissues. Quantification of HSP47 and Collagen I expression in stromal areas adjacent to KRT14+ squamous lesions was summarized in Tables S4-6. **(G)** Lung tissue specimens from patients with IPF were stained as in **(F)**. **(H)** Lung lesions with evidence of non-malignant squamous metaplasia were stained as in **(F).** Squamous lesions positive for KRT14 staining were identified. **(I)** Healthy bronchi from autopsied lungs stained as in **(F)** were used as negative controls.

**Supplementary Figure S7:**
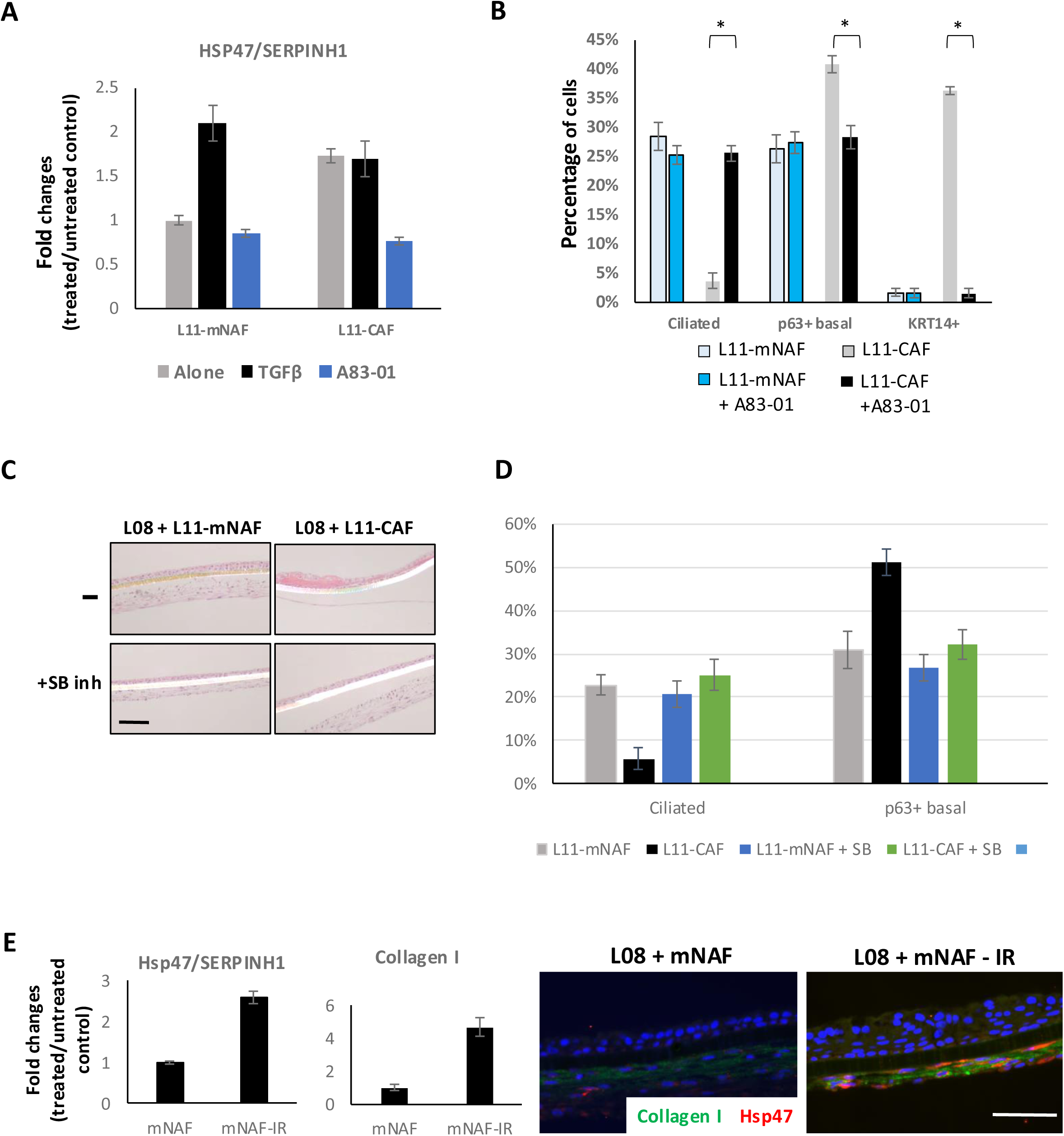
Fibroblasts Alone are SuLicient to Mediate Lung Squamous Metaplasia Through Activation of the TGFß Pathway and Downstream Engagement of HSP47. **(A)** mNAFs or CAFs were treated with TGF-β (10 ng/mL) or TGF-β inhibitor A83-01 (1 nM) for 48 h in 2D culture. HSP47 transcript expression was assessed by qPCR. Fold changes in mRNA expression are shown. **(B)** hBECs were co-cultured with mNAFs and CAFs under a double-sided condition in the presence or absence of A83-01 (1 nM) for 21 days. Quantification of cell lineage representation in the formed epithelium based on immunostaining is shown. **(C)** hBECs were co-cultured with mNAFs or CAFs under a double-sided condition in the presence or absence of the TGF-β pathway inhibitor SB431542 (10 nM) for 21 days. The formed epithelium was collected to assess morphology by H&E staining. **(D)** Quantification of cell lineage representation in hBECs cocultured with mNAFs or CAFs, in the presence or absence of the SB431542 inhibitor, based on immunostaining is shown. (**E)** The mRNA expression of Serpin H1/Hsp47 and Collagen I was analyzed by qPCR in irradiated or naïve mNAFs (left). Immunostaining showing expression of Hsp47 and Collagen I in cocultures with irradiated or naïve mNAFs (right).

**Supplementary Figure S8:**
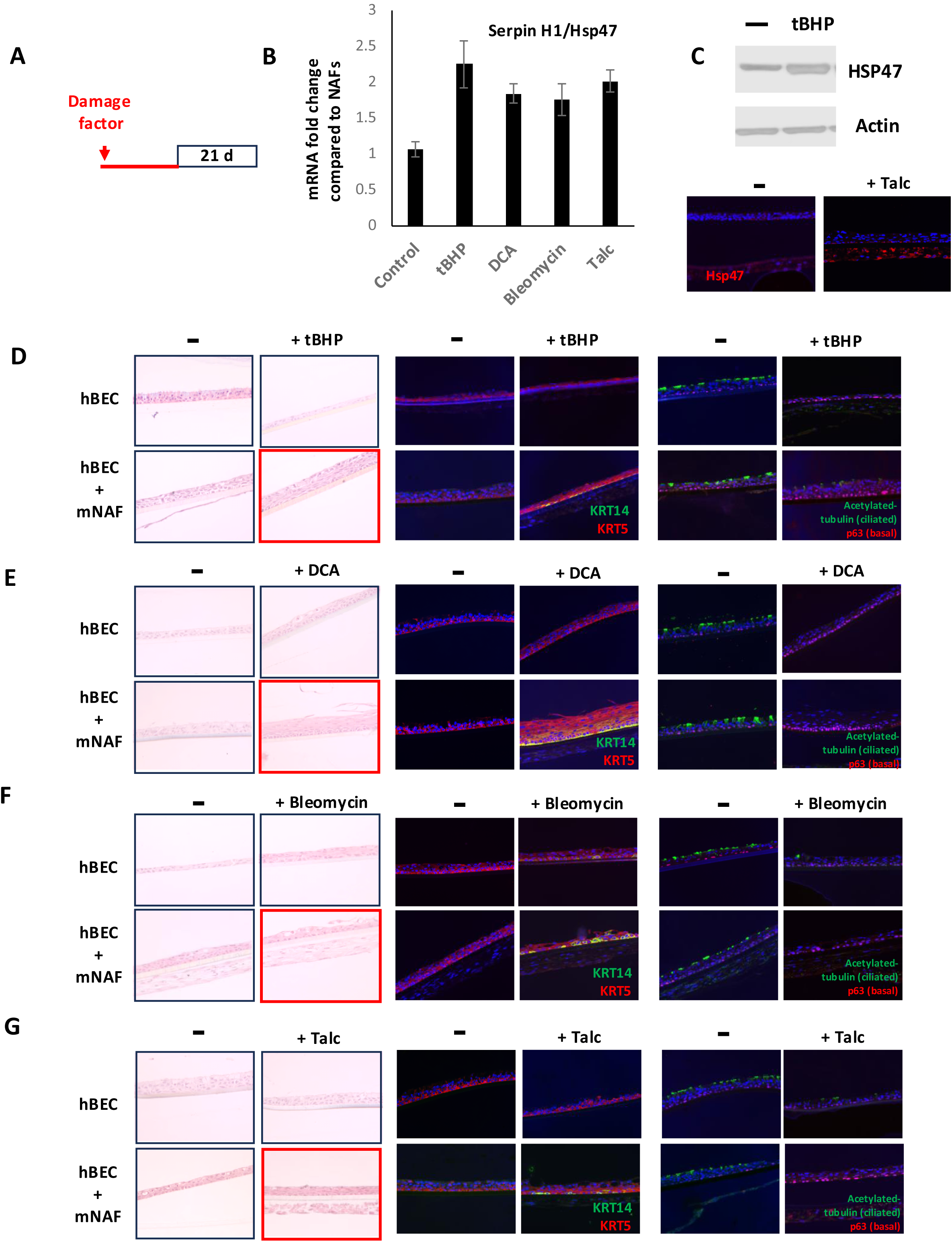
Stress Acts through Fibroblasts, not Epithelial Cells, to Induce Squamous Metaplasia in hBECs. Agents known to cause lung injury and increase risk for lung cancer (tert-butylhydroperoxide (tBHP)(an inducer of oxidative stress), Deoxycholic acid (DCA; bile salts), bleomycin (a therapeutic drug) and talc, each induced HSP47 expression in mNAFs at both mRNA and protein levels and when co-cultured with hBECs, squamous metaplasia was induced. **(A)** Schematic timeline showing the coculture procedure using fibroblasts pretreated with various damage agents or the vehicle control for 7 days. **(B)** The mRNA expression of Serpin H1/Hsp47 was analyzed by qPCR in fibroblasts treated with various damage agents or a vehicle control. **(C)** The protein expression of Hsp47 in mNAFs treated with 25 µM of tBHP or a vehicle control was compared by western blotting. Actin was used a loading control (top). The protein expression of Hsp47 was compared by immunostaining in cocultured mNAFs pretreated with 0.5 g/ml of talc powder or PBS (bottom). **(D-G)** Fibroblasts (mNAFs) were treated with a vehicle control or 25 µM tBHP **(D)** or 50 µM of DCA (Deoxycholic Acid) **(E)** or 250 µg/ml of Bleomycin **(F)** or 0.5 g/ml of Talc powder **(G)** for 7 days followed by coculture with hBECs for 21 days under ALI conditions. The formed epithelium was assessed for morphology by H&E staining and epithelial cell differentiation by immunostaining for bronchial epithelial and squamous metaplasia markers

**Supplementary Figure S9:**
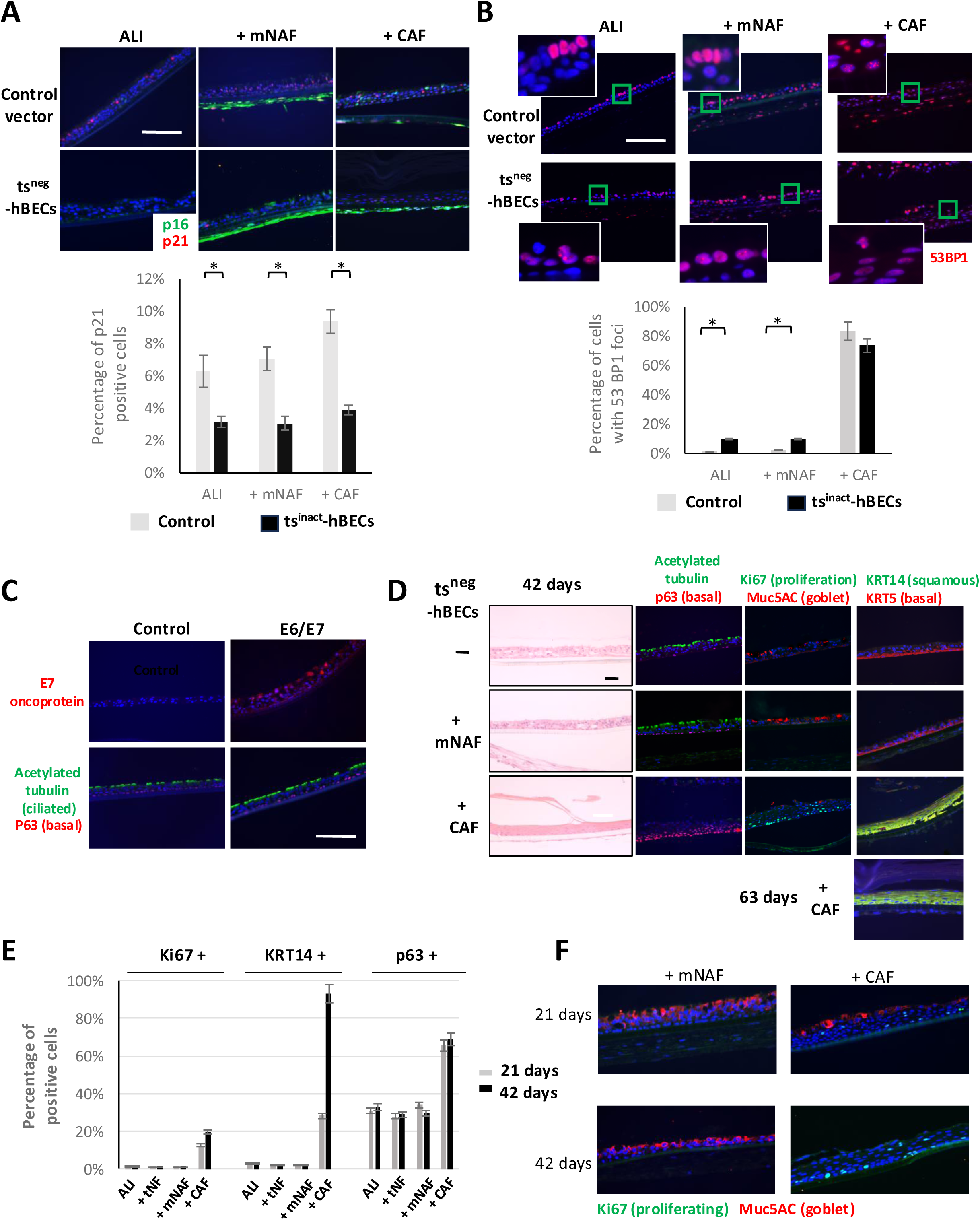
CAFs are Sufficient to Induce Squamous Dysplasia when Cocultured with hBECs Defective in p16 and p53 Pathways both *in vitro* and *in vivo.* **(A)** ts^neg^-hBECs or hBECs transduced with the control vector were co-cultured with or without fibroblasts. The expression of p16 and p21 was assessed using immunostaining (top). The number of p21-positive cells is plotted (bottom). There was no p16 expression in ts^neg^- hBECs under all conditions, but fibroblasts expressed high levels of p16. **(B)** ts^neg^-hBECs or hBECs transduced with the control vector were co-cultured with or without fibroblasts. The expression of 53BP1 was assessed by immunostaining. Magnified insets show an increase in nuclear 53BP1 foci in the squamous epithelium induced by co-culture with CAFs (top). Quantification of 53BP1 positive cells is shown (bottom). **(C)** ts^neg^-hBECs were cultured in ALI conditions for 21 days. The expression of E7 protein and a ciliated cell marker was assessed by immunostaining. **(D)** ts^neg^-hBECs were co-cultured with or without fibroblasts using a double-sided system for 42 or 63 days. The resulting epithelial structures were sectioned for H&E or immunofluorescence staining to assess the expression of specific markers. **(E)** Quantifications of Ki67+, KRT14+ and p63+ cells in ts^neg^-hBECs cultured in ALI or co-cultured with tNFs, mNAFs or CAFs for 21 days or 42 days are shown. **(F)** Bronchial lineage differentiation and proliferation were visualized by immunofluorescent staining for Muc5AC and Ki67 respectively in ts^neg^-hBECs cocultured with fibroblasts for 21 or 42 days.

**Supplementary Figure S10:**
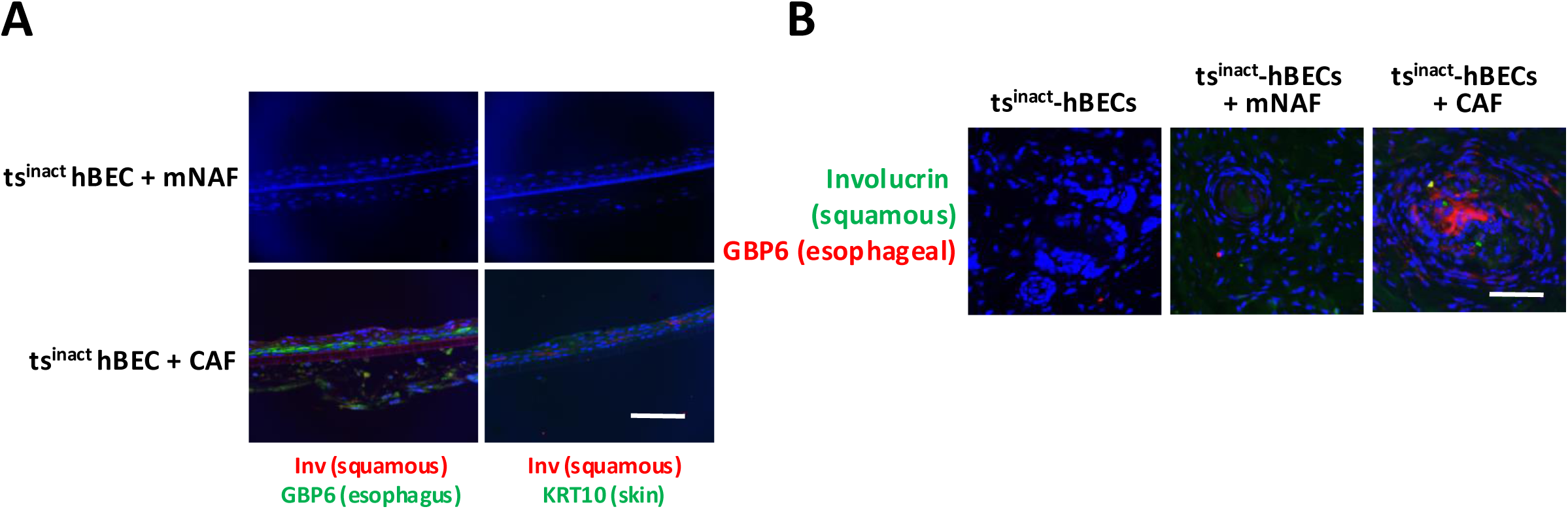
CAF-induced squamous dysplasia in both in vitro and in vivo recombinants expresses a marker of esophageal squamous epithelium. **(A)** Immunostaining-based characterization of esophageal-like squamous metaplasia in ts^neg^-hBECs cocultured with mNAFs or CAFs in ALI. **(B)** Immunostaining-based characterization of esophageal-like squamous metaplasia in ts^neg^-hBECs coinjected with mNAFs or CAFs in mice.

**Supplementary Figure S11:**
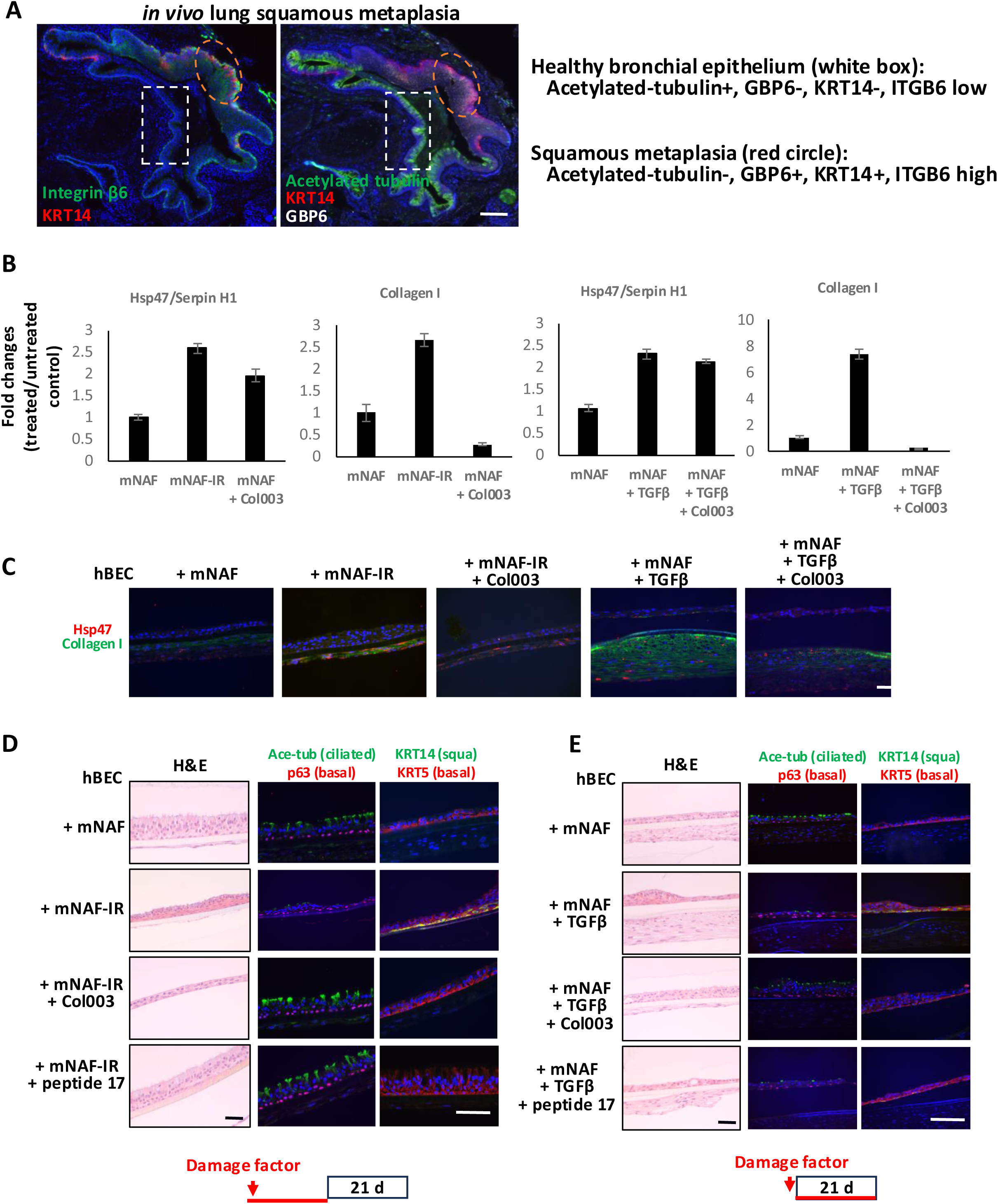
Clinical Specimens, as well as Irradiated and TGF-b-exposed Fibroblasts, show evidence that Squamous Metaplasia is Induced via HSP47/Integrin-mediated Mechanotransduction. In clinical metaplastic human lung specimens, ITGB6 expression was more intense and widespread in regions of squamous metaplasia where KRT14 and GBP6 expression was high, compared to regions of normal bronchial epithelium with high expression of ciliated cell markers. **(A)** ITGB6 and KRT14 expression was assessed by multiplex immunostaining of non-malignant lung squamous metaplasia specimens. The expression of GBP6 and acetylated tubulin was also evaluated in the same lesion, which showed areas of squamous metaplasia (orange oval) and neighboring columnar bronchial epithelium (white rectangle). **(B)** mNAFs were irradiated (20 Gy) following by treatment with10 μM of Col003 for 48 hrs. The mRNA expression of Serpin H1/Hsp47 or Collagen I was compared by qPCR in naïve, irradiated and Col003-treated cells (left). mNAFs were treated with 10 ng/ml of TGF-β or a vehicle control in the presence or absence of 10 μM of Col003 for 48 hrs. The mRNA expression of Serpin H1/Hsp47 or Collagen I was compared by qPCR in treated and untreated cells (right). **(C)** The protein expressions of Hsp47 and Collagen I were compared in cocultured fibroblasts treated various agents or a vehicle control by immunofluorescent staining. **(D)** Following the treatment procedure shown as a schematic timeline at the bottom, hBECs were cocultured for 21 days with naïve or irradiated mNAFs for 21 days in the presence or absence of 10 μM of Col003 or 2 mM of peptide 17. The formed epithelium was collected to assess morphology by H&E staining and epithelial cell differentiation status by immunostaining for the expression of bronchial epithelial markers and a squamous marker. **(E)** Following the treatment procedure showed as a schematic image on the bottom, hBECs were coculture with TGF-β- or vehicle control-treated fibroblasts (mNAFs) for 21 days in the presence or absence of 10 μM of Col003 or 2 mM of peptide 17. The formed epithelium was collected to assess morphology by H&E staining and epithelial cell differentiation status by immunofluorescence staining for the expression of bronchial epithelial markers and a squamous marker.

**Supplementary Figure S12:**
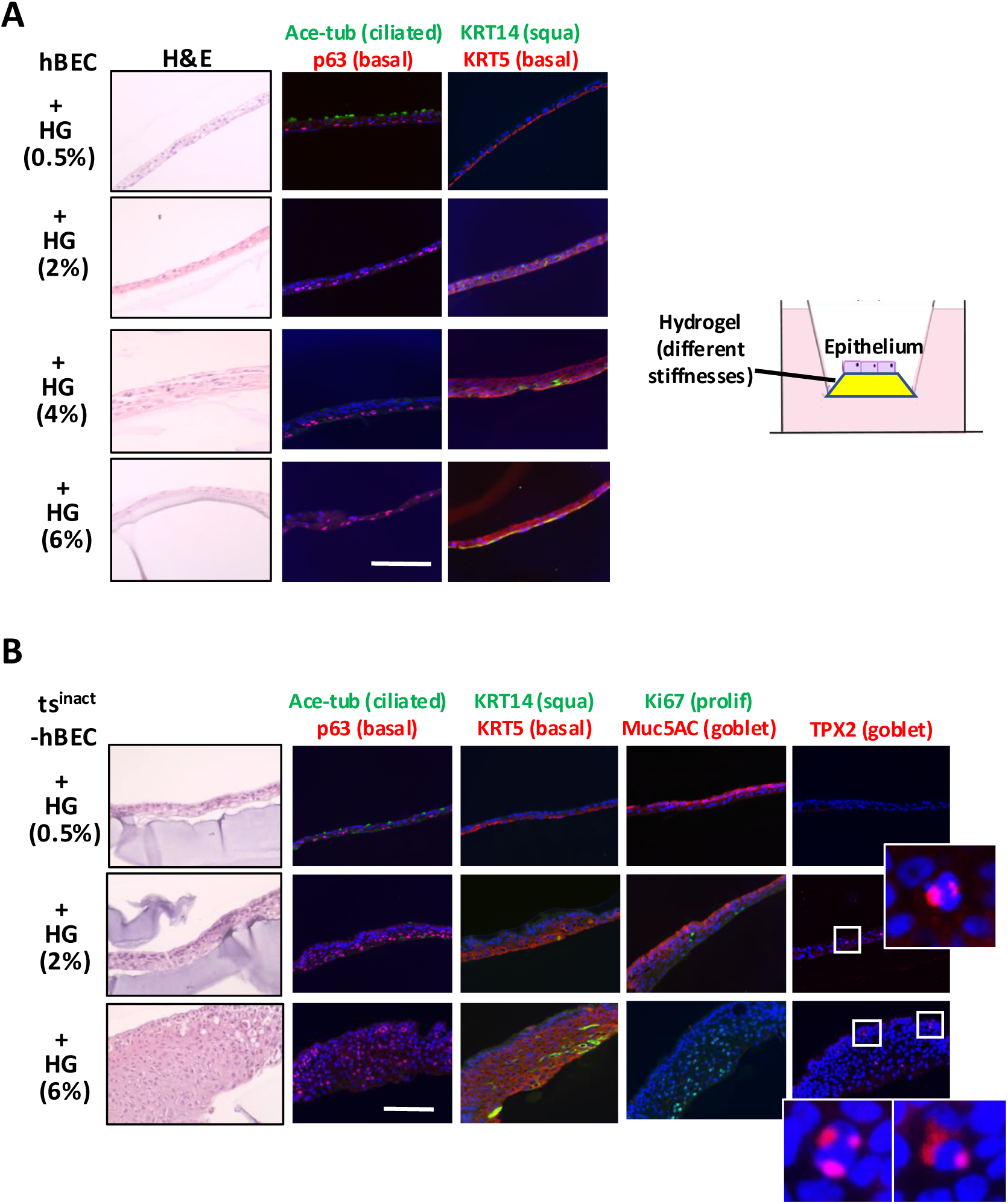
StiLer Matrix Can Directly Control the Progression of Squamous Metaplasia and Dysplasia Through Activation of Mechanotransduction. **(A)** Schematic representation of epithelial cells grown on different concentrations of hydrogel for 14 days in ALI (right). hBECs cultured on different concentrations of hydrogel were assessed for morphology by H&E staining and epithelial cell differentiation by immunostaining for selected markers (left). **(B)** ts^neg^-hBECs were cultured on different concentrations of hydrogel for 14 days, and morphology was assessed by H&E staining and epithelial cell differentiation, proliferation, and aneuploidy by immunostaining of selected markers. Magnified insets show dividing cells with three spindle poles marked by TPX2 staining.

**Supplementary Figure S13:**
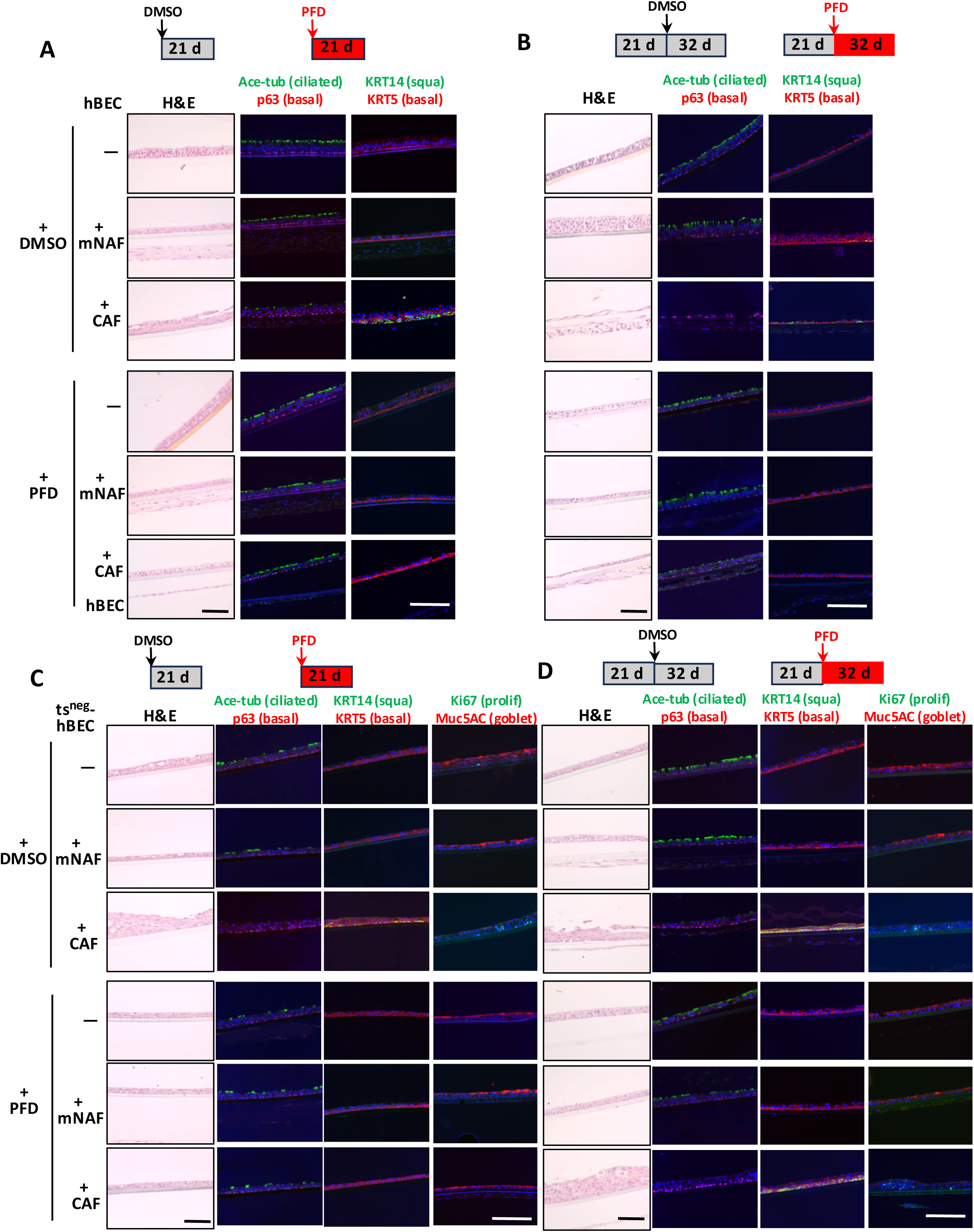
Blocking Hsp47-dependent Pathways Effectively Prevents and Reverts Fibroblast-induced Squamous Metaplasia but only Prevents, Not Reverts, Fibroblast-induced Dysplasia. Effects of the anti-fibrotic agent Pirfenidone (PFD) were evaluated by H&E staining and immunostaining for bronchial differentiation (ciliate and goblet) and squamous (basal) markers in various experimental set ups illustrated at the top of each panel. **(A)** hBECs cultured alone or with mNAFs or CAFs treated with PFD or vehicle control for 21 days prior to assessment. **(B)** hBECs cultured alone or with mNAFs or CAFs for 21 days followed by treatment with PFD or vehicle control for 32 days prior to assessment. **(C)** ts^neg^-hBECs cultured alone or with mNAFs or CAFs treated with PFD or vehicle control for 21 days prior to assessment. **(D)** ts^neg^-hBECs cultured alone or with mNAFs or CAFs for 21 days followed by treatment with PFD or vehicle control for 32 days prior to assessment.

**Supplementary Figure S14:**
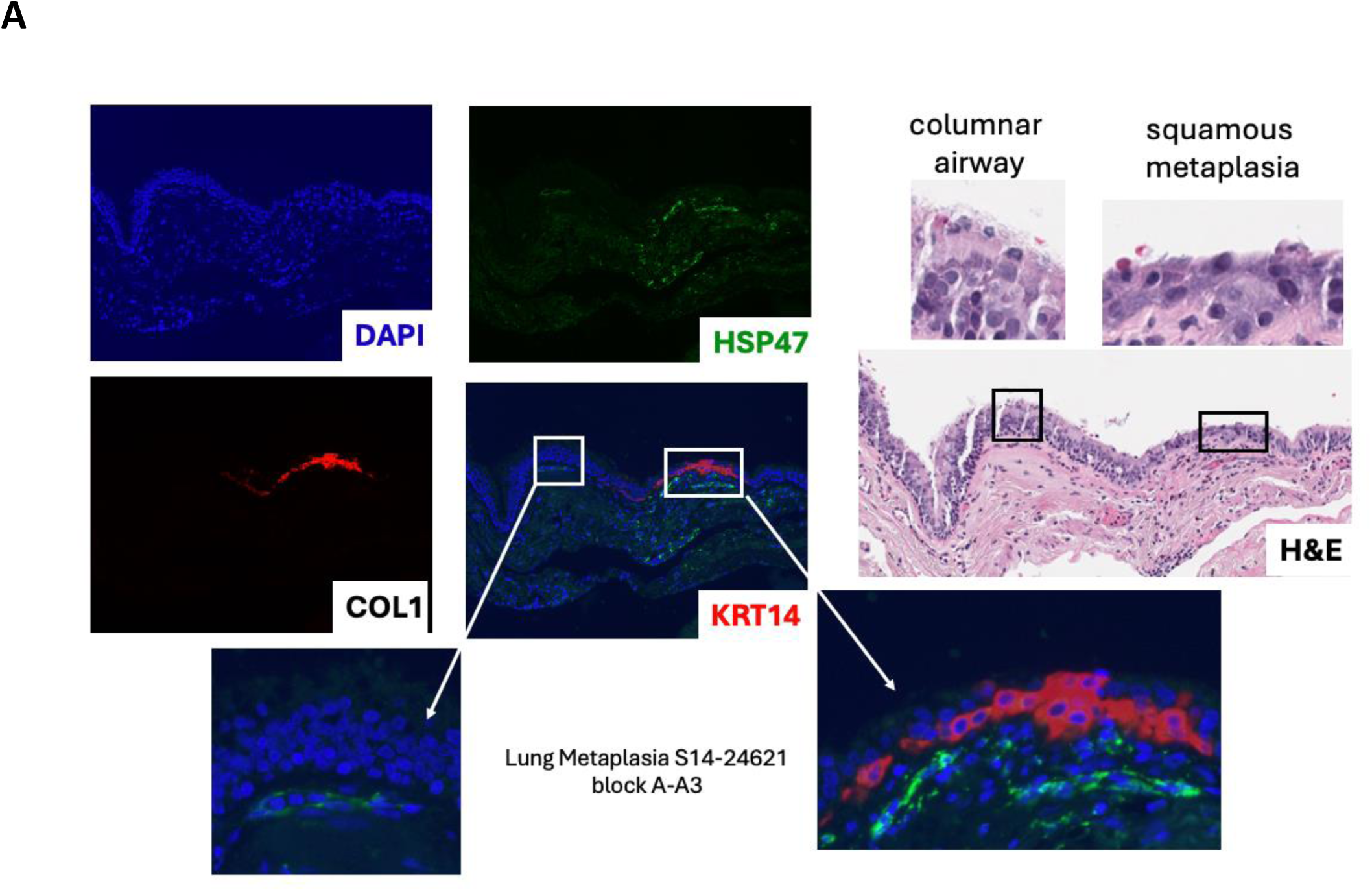
Patterns of Tissue Stress Response and Focal Consequences in Injured Bronchial Tissue. **(A)** Non-malignant lung squamous metaplasia stained for HSP47 and KRT14 showing concomitant presence of KRT14-positive squamous lesions associated with underlying foci of HSp47 expression (right section of the immunostaining image) and focal areas with high HSP47 expression without associated KRT14-positive squamous lesion (left section of the immunostaining image). Magnified insets illustrate these two types of patterns. The region with pseudostratified epithelium or squamous metaplasia were identified in H&E staining in the corresponding tissue (right)

**Supplementary Figure S15:**
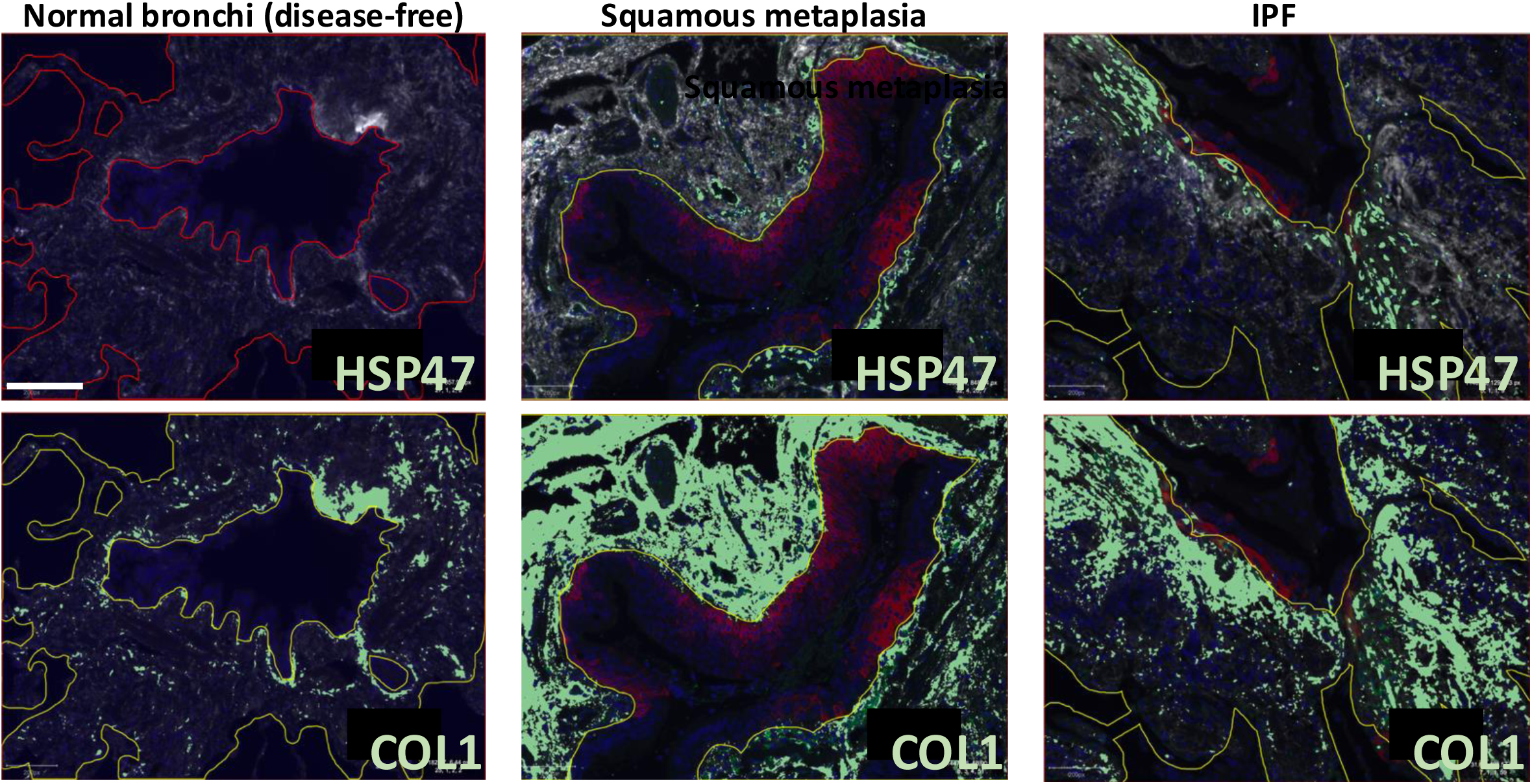
Quantitative analysis of HSP47 and Collagen 1 expression in healthy and diseased lung. Representative images of mIHC for healthy lung (left column), non-malignant lung squamous metaplasia (middle column), and fibrotic lung (Idiopathic Pulmonary Fibrosis (IPF); right column). Pixels positive above the threshold were masked (green color), and positivity was expressed as the percentage of masked area/total stromal area (see Tables S4-S6).

**Supplementary Table S1:** Summary of epithelial phenotypes observed when bronchial epithelial cells, isolated from matched histologically normal lung specimens, were co-cultured with mNAFs and CAFs derived from different LSCC tumor specimens under the double-sided condition.

| Co-culture | L08-hBEC | L10-hBEC | L11-hBEC | Total events<br>(Pseudo-stratified) | Total events<br>(Squamous) |
| --- | --- | --- | --- | --- | --- |
| **mNAF**<br>**L09** | Pseudostratified<br>(n=2/2) | Pseudostratified<br>(n=2/2) | Pseudostratified<br>(n=2/2) | **6/6** |  |
| **L10** | Pseudostratified<br>(n=4/4) | Pseudostratified<br>(n=2/2) | Pseudostratified<br>(n=2/2) | **8/8** |  |
| **L11** | Pseudostratified<br>(n=38/42) | Pseudostratified<br>(n=4/5) | Pseudostratified<br>(n=3/3) | **45/50** |  |
| **L14** | Pseudostratified<br>(n=1/4) | Pseudostratified<br>(n=2/2) | N/A | **3/6** |  |
| **L21** | Pseudostratified<br>(n=4/4) | Pseudostratified<br>(n=2/2) | N/A | **6/6** |  |
| **CAF**<br>**L09** | Pseudostratified<br>(n=2/2) | Pseudostratified<br>(n=2/2) | Squamous<br>(n=2/2) |  | **2/6** |
| **L10** | Squamous<br>(n=3/4) | Squamous<br>(n=1/2) | Squamous<br>(n=2/2) |  | **6/8** |
| **L11** | Squamous<br>(n=46/50)* | Squamous<br>(n=4/5) | Squamous<br>(n=3/3) |  | **53/58** |
| **L14** | Squamous<br>(n=8/8) | Squamous<br>(n=2/2) | N/A |  | **10/10** |
| **L21** | Squamous<br>(n=4/4) | Squamous<br>(n=2/2) | N/A |  | **6/6** |

- **Pseudostratified:** A pseudostratified bronchial epithelial structure is defined as exhibiting beating cilia, as well as expression of cilia proteins (Acetylated gamma-tubulin), club cell secretions (CCL10), goblet cell secretions (MUC5AC) and basal cell markers (p63). Additionally, <5% of basal cells express KRT14 based on immunofluorescent staining - **Squamous Metaplasia:** A squamous metaplastic structure is defined as being devoid of cilia, and lacking expression of club cell and goblet cell proteins plus >5% of basal cells express KRT14, a squamous marker, based on the immunofluorescent staining
\*(n=46/50) means that of the 50 times that these two cell populations were cultured together in ALI, 46 times they generated a squamous metaplastic structure in greater than 5% of the cultured area and 4 times they did not.

**Supplementary Table S2:**
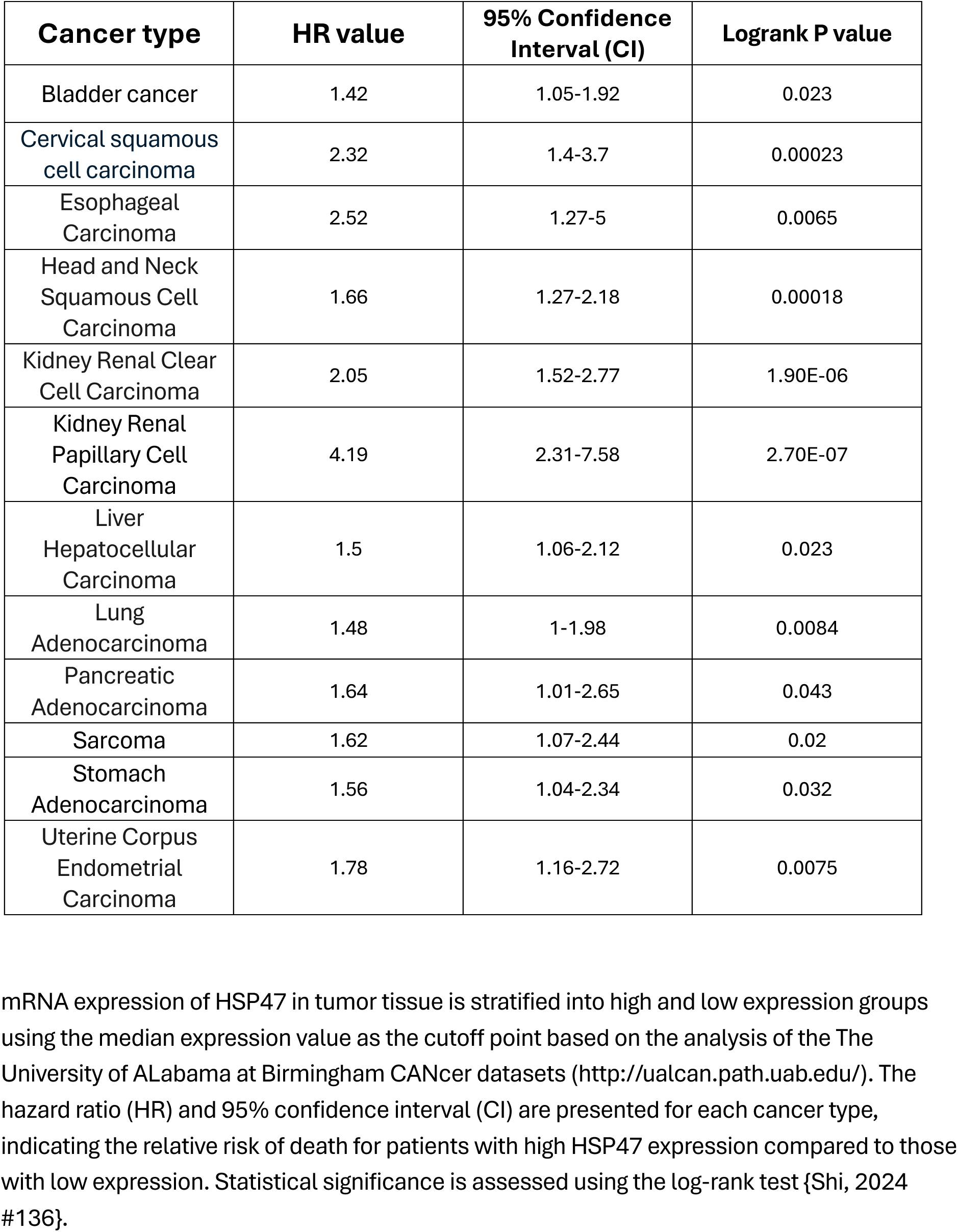
The relative risk of death for patients with high HSP47 expression compared to those with low expression (modified from Shi, et al., 2024).

| **Cancer type** | **HR value** | **95% Confidence Interval (CI)** | **Logrank P value** |
| --- | --- | --- | --- |
| Bladder cancer | 1.42 | 1.05-1.92 | 0.023 |
| Cervical squamous cell carcinoma | 2.32 | 1.4-3.7 | 0.00023 |
| Esophageal Carcinoma | 2.52 | 1.27-5 | 0.0065 |
| Head and Neck Squamous Cell Carcinoma | 1.66 | 1.27-2.18 | 0.00018 |
| Kidney Renal Clear Cell Carcinoma | 2.05 | 1.52-2.77 | 1.90E-06 |
| Kidney Renal Papillary Cell Carcinoma | 4.19 | 2.31-7.58 | 2.70E-07 |
| Liver Hepatocellular Carcinoma | 1.5 | 1.06-2.12 | 0.023 |
| Lung Adenocarcinoma | 1.48 | 1-1.98 | 0.0084 |
| Pancreatic Adenocarcinoma | 1.64 | 1.01-2.65 | 0.043 |
| Sarcoma | 1.62 | 1.07-2.44 | 0.02 |
| Stomach Adenocarcinoma | 1.56 | 1.04-2.34 | 0.032 |
| Uterine Corpus Endometrial Carcinoma | 1.78 | 1.16-2.72 | 0.0075 |

mRNA expression of HSP47 in tumor tissue is stratified into high and low expression groups using the median expression value as the cutoff point based on the analysis of the The University of ALabama at Birmingham CANcer datasets (http://ualcan.path.uab.edu/). The hazard ratio (HR) and 95% confidence interval (CI) are presented for each cancer type, indicating the relative risk of death for patients with high HSP47 expression compared to those with low expression. Statistical significance is assessed using the log-rank test {Shi, 2024 #136}.

**Supplementary Table S3:** Patient diagnosis for tissue specimens used in multiplex immunofluorescent staining.

| Diagnosis | Patient-tissue block | Stroma adjacent to KRT14+ squamous lesions |  |  |  |
| --- | --- | --- | --- | --- | --- |
|  |  | Hsp47+Col1high | Hsp47+Col1low | Hsp47-Col1high | Hsp47-Col1low |
| Lung metaplasia | LM01-1 | 6 | 1 |  |  |
|  | LM01-2 | 2 |  | 2 |  |
|  | LM01-3 | 3 | 1 | 1 |  |
|  | LM02-1 | 6 |  | 1 |  |
|  | LM03-1 | 2 |  | 2 | 1 |
|  | LM04-1 |  | 5 |  |  |
|  | LM04-2 | 4 | 6 |  |  |
|  | LM05-1 | 6 |  |  |  |
|  | LM05-2 | 5 |  |  |  |
|  | LM05-3 | 7 | 1 |  |  |
|  | LM05-4 |  | 2 | 2 |  |
|  | LM06-1 | 4 |  | 1 | 1 |
|  | LM07-1 | 2 |  |  |  |
|  | LM07-2 | 4 |  |  |  |
|  | LM07-3 | 5 | 1 |  |  |
|  | LM07-4 | 5 |  |  |  |
|  | LM07-5 | 3 |  | 1 |  |
|  | LM08-1 | 5 |  | 1 |  |
|  | LM09-1 |  |  |  | 3 |
| LSCC | LSCC1 | 3 | 2 |  |  |
|  | L08 | 3 | 2 |  |  |
|  | L10 | 1 |  | 1 |  |
|  | L11 | 7 | 1 |  |  |
|  | L21 | 4 | 1 |  |  |
| Number of images |  | 87 | 23 | 12 | 5 |
| Hsp47/Col1 expression profile in stroma adjacent o KRT14+ lesions (% of images) |  | 68.5% | 18.11% | 9.45% | 3.94% |

**Supplementary Table S4:** Quantification of HSP47 and Collagen I expression in stromal areas adjacent to KRT14+ squamous lesions in LSCC or lung squamous metaplasia, or in stromal areas adjacent to airway epithelial structures in disease-free (true normal) or matched normal lung tissues.

| **Tissue types** | **Tissue resource** | **KRT14+** | **Hsp47-Col1low** | **Hsp47-Col1high** | **Hsp47+ Col1high** | **Hsp47+ Col1low** |
| --- | --- | --- | --- | --- | --- | --- |
| **Disease-free lung** | **Autopsy tNL01** | 0 | 9 |  |  |  |
| **Matched normal lung** | Adjacent to LSCC1 | 0 | 5 |  |  |  |
|  | Adjacent to LSCC-L08 | 0 | 3 | 1 | 1 |  |
|  | Adjacent to LSCC-L10 | 0 | 1 | 3 | 1 |  |
|  | Adjacent to LSCC-L11 | 0 | 1 | 4 |  |  |
|  | Adjacent to LSCC-L21 | 0 | 1 | 5 |  |  |
| **Number of images** |  | 0 | 20 | 13 | 2 |  |
| **Hsp47/Col1 expression profile in stroma (% of images)** |  | **0** | **57.14%** | **37.14%** | **5.71%** | **0** |

**Supplementary Table S5:** Summary of Table S3 and S4 showing the distribution of KRT14-positive lesions based on HSP47 and Collagen I expression status in 5 LSCC and 20 lung squamous metaplasia specimens collected from 10 patients without LSCC.

**Supplementary Table S6:** Summary of Table S3 and S4 showing lack of KRT14 expression and low HSP47 expression in disease-free or matched normal lungs.

